# Diversifying particle-orientation distributions in cryo-EM with LEA protein additives

**DOI:** 10.64898/2026.07.09.737610

**Authors:** Kaitlyn M. Abe, Timothy Grant, Ci Ji Lim

## Abstract

Preferential orientation at the air-water interface (AWI) remains a persistent challenge in cryo-electron microscopy, often requiring extensive optimization or specialized grid preparation strategies. Here, we show that late embryogenesis abundant (LEA) proteins, previously shown to protect fragile samples from AWI-induced damage, can be leveraged to resolve sample orientation bias. The addition of LEA proteins can yield either new orientation distributions or more isotropic ones in samples suffering from orientation biases. These findings establish LEA proteins as a practical class of cryo-EM additives that can reduce AWI-induced damage and improve particle-orientation distributions in single-particle analysis.

## Introduction

Cryogenic electron microscopy (cryo-EM) has become a cornerstone method in structural biology. Advances in microscopes, detectors, image processing, and reconstruction algorithms have made it possible to determine high-resolution structures of macromolecular complexes, including some that were previously inaccessible by other methods^1,2^, yet cryo-EM sample preparation remains a bottleneck. In standard single-particle analysis (SPA), particles are applied to a grid, blotted into a thin aqueous film, and rapidly plunge-frozen. During this process, particles are exposed to the air-water interface (AWI), which can disrupt complexes, promote aggregation or denature fragile particles^3–5^. Even when the particles survive this exposure, they almost always adsorb to the AWI^6^, often adopting only a limited set of views. Severe preferred orientation can lead to insufficient angular sampling and produces anisotropic three-dimensional (3D) reconstructions with poorly resolved features^2,7–9^. To overcome preferred orientation issues, researchers utilized additives such as detergents^10,11^, engineered polymers^12^, or applied support films^13^. However, these strategies are often complex to deploy, could lead to artificial interactions between the additives and sample^14^, and require iterative optimization.

We previously showed that Late Embryogenesis Abundant (LEA) proteins can mitigate AWI-induced sample damage during cryo-EM grid preparation^15^. LEA proteins are small, highly soluble proteins associated with desiccation tolerance in plants^16,17^ and other stress-tolerant organisms, such as nematodes^18–21^, tardigrades^22^, and insects^23^. Although many LEA proteins are intrinsically disordered in solution, they are predicted to adopt an α-helical structure at air-water interfaces that confers specific adsorption affinity to the interface^24–26^. This unique feature presumably led to their protective function during cryo-EM sample plunge freezing.

In our previous work, the two tested LEA proteins, AavLEA1 and RvLEAM_short_, did not produce a uniform effect across all samples. Instead, the two proteins led to different orientation distributions for the same sample, and the distributions are seemingly sample-dependent^15^. This behavior suggests some degree of interaction between LEA proteins and sample particles; however, we did not observe additional structured density in the reconstructed cryo-EM maps that would indicate a stable, specific interaction.

Here, we tested whether this LEA-dependent reshaping of particle orientations could be exploited as a strategy to improve cryo-EM orientation distributions. We first examined catalase, a benchmark sample that commonly adopts a strongly preferred particle orientation^10,11^. Adding LEA proteins AavLEA1 or RvLEAM_short_ produced orientation distributions that were orthogonal and thus complementary to the control. Consequently, combining them led to a more complete orientation distribution, and improved map anisotropy metrics. We next tested hemagglutinin (HA) trimer, which exhibits a more severe orientation bias^8,27,28^, and requires stage-tilt data collection to gain additional views for an isotropic reconstruction^8^. Adding LEA proteins yielded a significantly broader orientation distribution than the control, enabling isotropic, high-resolution maps of the HA trimer without relying on stage tilting during data collection.

To test whether LEA-dependent orientation control could be expanded beyond the two tested LEA proteins, we identified four additional LEA proteins and tested their effects on the human Polα-primase complex, which suffers from AWI-induced damage^15^. All four new proteins mitigated AWI-induced damage to the complex. More importantly, all six LEA proteins, including AavLEA1 and RvLEAM_short_^15^, produced a distinct orientation distribution. These results thus demonstrate that LEA proteins can be used not only to protect cryo-EM samples but also diversify particle orientations. Together, this work presents a new strategy for overcoming particle-orientation bias in cryo-EM single particle analysis. By screening a catalog of LEA protein additives, users can identify conditions that preserve sample integrity and produce more complete orientation distributions; selected LEA conditions can then be used individually or combined during data processing to enable more isotropic 3D reconstructions.

## Results

### LEA proteins alter particle orientation of human catalase

Catalase particles adopt a dominant orientation at the AWI on cryo-EM grids^10–13^, making catalase an ideal test specimen for determining whether AavLEA1 or RvLEAM_short_ can alter particle-orientation distributions. We found that both proteins significantly changed the catalase particle orientation distribution (**Fig. 1, Supplementary Figs. 1-3**). Two-dimensional (2D) classification showed that catalase particles in both LEA datasets predominantly adopted a “side-view”, whereas the control dataset particles were mostly “front-view” (**Fig. 1a**). However, the two LEA proteins did not produce identical distributions: compared with AavLEA1, RvLEAM_short_ retained a higher proportion of front views, yielding a slightly more complete orientation distribution (**Fig. 1a, right**).

**Figure 1:**
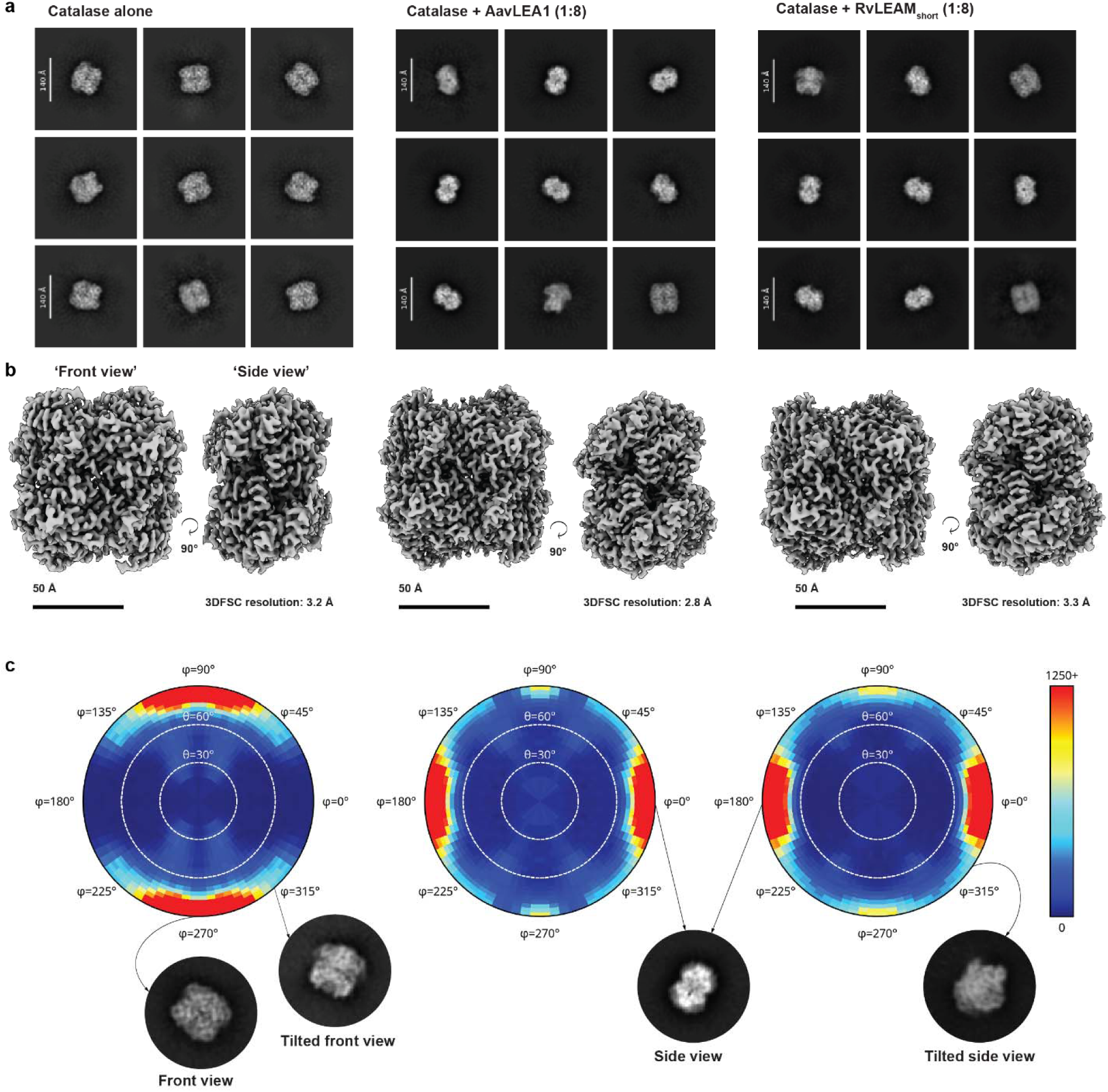
LEA proteins induce an orientation shift in human erythrocyte catalase. **(a)** Representative 2D classes of the final particle stacks from the catalase alone (left), catalase + AavLEA1 (middle), and catalase + RvLEAM_short_ (right). **(b)** Cryo-EM map of catalase showing top and side views with catalase control, catalase + AavLEA1 and catalase + RvLEAM_short_ from left to right. **(c)** Orientation distribution diagram of each dataset showing select 2D classes from different points on the diagram with catalase control, catalase + AavLEA1 and catalase + RvLEAM_short_ from left to right.

To fairly compare reconstructed 3D map quality across datasets, we processed the datasets using the same workflow and reconstructed their final cryo-EM maps using the same number of particles and initial reference volume. Although their global map resolutions were comparable at ∼3 Å, as assessed by 3DFSC^8,29^ (**Fig. 1b, Supplementary Figs. 1-3**), their orientation distributions differed substantially (**Fig. 1c**). The control exhibited a narrow top-bottom preferred orientation, whereas both LEA-datasets displayed orthogonal but broader distributions (**Fig. 1c**). To quantify these differences, we used isotropy metrics; Sample Compensation Factor (SCF)^30,31^, conical FSC Area Ratio (cFAR), and sphericity^8,30^. The control map was least isotropic (SCF = 0.28, cFAR = 0.23, sphericity = 0.93), while the LEA proteins improved isotropy: AavLEA1 yielded an SCF of 0.63, cFAR of 0.49, and sphericity of 0.96, and RvLEAM_short_ led to an SCF of 0.58, cFAR of 0.59, and sphericity of 0.96 (**Supplementary Table 1**).

Because the LEA-datasets particles had orthogonal views to that of the control dataset, we tested whether combining two datasets would yield a more comprehensive orientation distribution and improve map quality. We selected the AavLEA1 dataset for combination with the control dataset because they showed the most orthogonal distribution relative to each other. An equal number of particles were used from each dataset. The merged particle stack gave a substantially more balanced distribution, increasing the values of SCF to 0.80, cFAR to 0.70, and sphericity to 0.97 (**Supplementary Fig. 4**). These improvements demonstrate that LEA-induced orientation redistribution can be exploited computationally by combining datasets with complementary views. Although all the reconstructions were of sufficient quality for accurate model building, anisotropic resolution is visible in the catalase control map as apparent stretching in the front view and in the LEA protein maps as apparent stretching in the side view. Neither are visible in the combined control + AavLEA1 map which produced the best density overall. Thus, catalase establishes the proof-of-concept that LEA proteins can reshape biased orientation distributions, paving a strategy to combine datasets with complementary orientation distributions to achieve better map isotropy.

### LEA protein addition led to an isotropic HA trimer cryo-EM map

HA trimer provides a more stringent test than catalase because its preferred orientation compromises 3D reconstruction. HA trimer particles adsorb predominantly in a “head-down” orientation at the AWI, a bias severe enough that prior work required stage-tilt data collection to obtain an isotropic 3D map^8,32,33^. This behavior has made HA trimer a gold-standard benchmark for testing orientation-bias mitigation strategies^4,8,12,14,17–21^. We therefore asked whether AavLEA1 or RvLEAM_short_ could improve HA trimer particle orientations sufficiently to support isotropic 3D reconstruction. Because the large contrast difference between head-down and side-view HA trimer particles can bias particle picking toward the brighter head-down views, we used denoised micrographs for all HA trimer datasets to minimize orientation bias during particle selection (see Methods; **Supplementary Figs. 5-7**).

The no-additive control contained primarily top-view 2D class averages and exhibited only ∼30° variation in angular distribution. In contrast, the addition of AavLEA1 or RvLEAM_short_ increased the angular coverage to approximately ∼60-70° from the top view, producing 2D classes that extended nearly to a full side-view orientation (**Fig. 2a**). To ensure a fair comparison of the reconstructed cryo-EM maps, we used the same number of particles from each dataset and the same initial reference volume. All three datasets yielded maps with comparable 3DFSC^8^-reported global resolutions of ∼3.5-3.7 Å (**Fig. 2b**). As expected, the map from the control dataset showed substantial anisotropy as compared to the AavLEA1 and RvLEAM_short_ datasets (**Fig 2c**). Isotropy metrics confirmed that the control dataset produced the most anisotropic map, with suboptimal SCF and cFAR values, and reduced sphericity (SCF = 0.30, cFAR = 0.03, sphericity = 0.83). In contrast, both LEA-containing datasets yielded substantially more isotropic reconstructions: AavLEA1 improved the scores to SCF = 0.92, cFAR = 0.55, and sphericity = 0.98, and RvLEAM_short_ improved the scores to SCF = 0.84, cFAR = 0.51, and sphericity = 0.98 (**Supplementary Table 2**).

**Figure 2:**
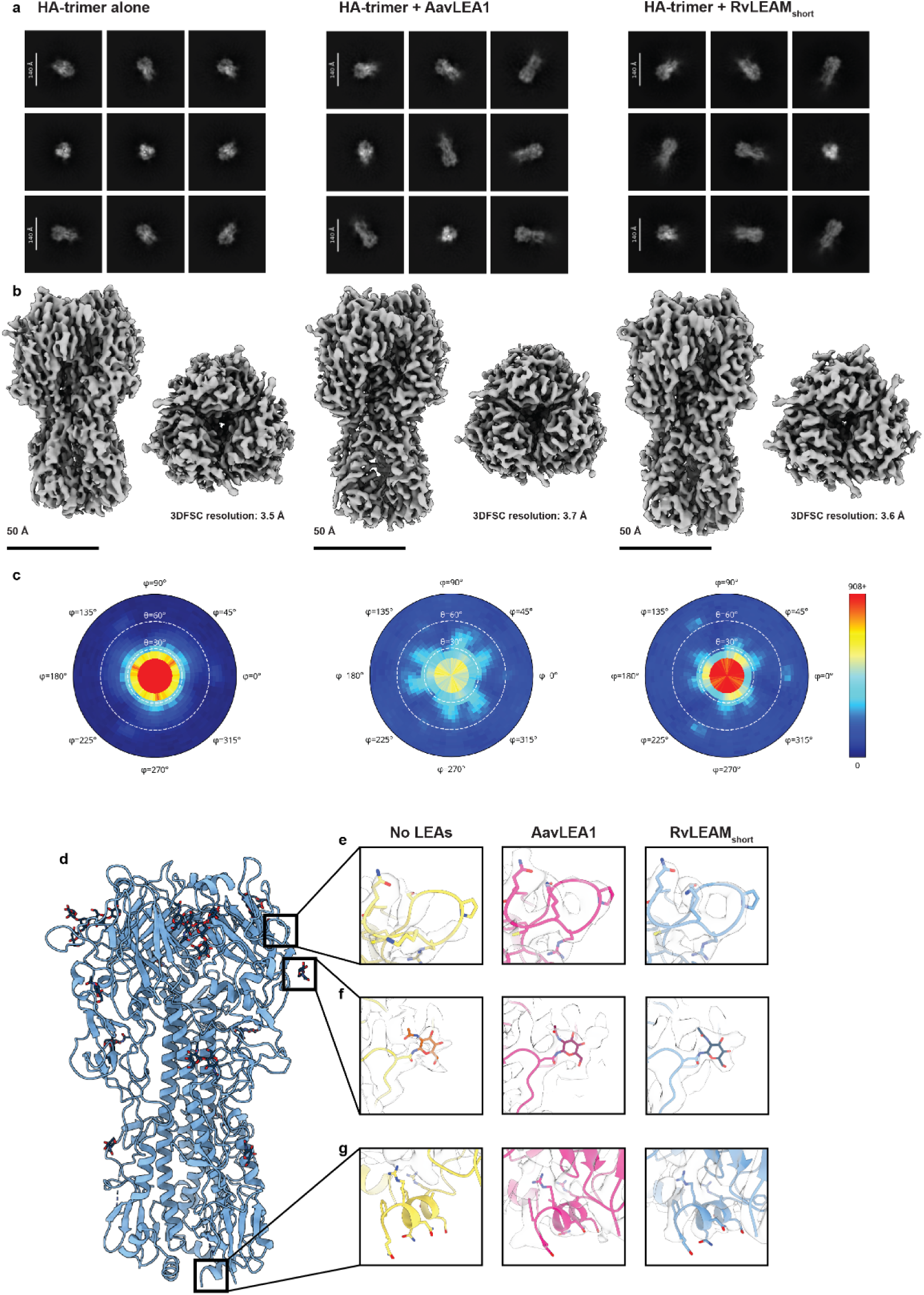
LEA proteins improve map isotropy for HA trimer. **(a)** Representative 2D classes of the final particle stacks from HA trimer alone (left), HA trimer + AavLEA1 (middle), and HA trimer + RvLEAM_short_ (right). **(b)** Cryo-EM map of HA trimer showing side and top views with HA trimer alone (left), HA trimer + AavLEA1 (middle), and HA trimer + RvLEAM_short_ (right). **(c)** Orientation distribution diagram of each dataset from left to right, HA trimer alone, HA trimer + AavLEA1, and HA trimer + RvLEAM_short_. **(d)** HA trimer model from RvLEAM_short_ map **(e)** Model-to-map fits for a representative loop region with residues 139-147. **(f)** Model-to-map fits for representative glycosylated residue 81 and corresponding glycan 609. **(g)** Model-to-map fits for C-terminal α-helix, residues 492-501.

These improvements in map isotropy were also reflected in map interpretability for atomic model building (**Fig. 2d-g**). Average Q-scores increased from 0.52 in the no-additive control to 0.64 and 0.66 in the AavLEA1 and RvLEAM_short_ reconstructions, respectively, indicating improved local resolvability despite comparable global resolutions. The improvement extended to glycosylation sites, which are often difficult to resolve because they are flexible and solvent-exposed^34^. Averaged across all modeled glycans, Q-scores increased from 0.48 in the control to 0.57 with AavLEA1 and 0.58 with RvLEAM_short_. The corresponding glycosylated residues showed a similar improvement, increasing from 0.53 in the control to 0.68 and 0.69 with AavLEA1 and RvLEAM_short_, respectively. These trends were also apparent at individual sites. At N81, the residue Q-score increased from 0.57 in the control to 0.74 with AavLEA1 and 0.68 with RvLEAM_short_, accompanied by improved density for the associated glycan, whose Q-score increased from 0.55 to 0.65 and 0.71, respectively (**Fig. 2f**). Together, these results show that LEA proteins improve map quality for flexible, solvent-exposed features, enabling more reliable modeling of glycosylated regions.

In addition, poorly resolved regions of the HA trimer also benefited from the addition of AavLEA1 or RvLEAM_short_. In the control reconstruction, the C-terminal domain exhibited weak density, with the C-terminal α-helix showing a low Q-score of 0.08 (**Fig. 2g, left**). In contrast, this region became well resolved in the presence of AavLEA1 and RvLEAM_short_, with Q-scores increasing to 0.51 and 0.56, respectively (**Fig. 2g, middle, right**). Similarly, the loop spanning residues 129–147 was less well defined in the control map, with an average Q-score 0.56 in this region, but showed improved density with both LEA proteins, reaching average Q-scores of 0.64 and 0.66, respectively (**Fig. 2e**). These site-specific improvements show that LEA proteins enhance map interpretability across multiple structurally challenging regions, even when global resolution changes little.

The better-resolved glycan density in the LEA-datasets may reflect more than improved orientation distribution. Because glycans are flexible and solvent-exposed, their improved density is also consistent with better preservation of local structure through LEA-mediated protection from AWI-induced damage. Thus, LEA proteins may improve not only map isotropy, but also the interpretability of local features that are otherwise difficult to model.

### An expanded LEA protein panel reveals broad AWI protection and distinct particle-orientation distributions

The catalase and HA trimer experiments showed that AavLEA1 and RvLEAM_short_ can reshape particle-orientation distributions, but their effects were sample- and LEA-dependent. This suggested that expanding beyond these two group 3 LEA proteins could provide a more diverse set of additives for mitigating preferred-orientation bias. We therefore expanded the LEA protein toolkit by testing four additional LEA proteins spanning groups 2, 3, and 4. These proteins were selected based on their molecular weight, reported roles in desiccation tolerance, and sequence features. The panel included the group 2 protein CuLEA from *Citrus unshiu*^35^(**Fig. 3a**), the group 3 proteins LEA7 from *Arabidopsis thaliana*^36,37^ and PvLEA4 from *Polypedilum vanderplanki*^38,39^ (**Fig. 3d-e**), and group 4 protein AtLEA4-5 from *A. thaliana*^40^ (**Fig. 3f**).

**Figure 3:**
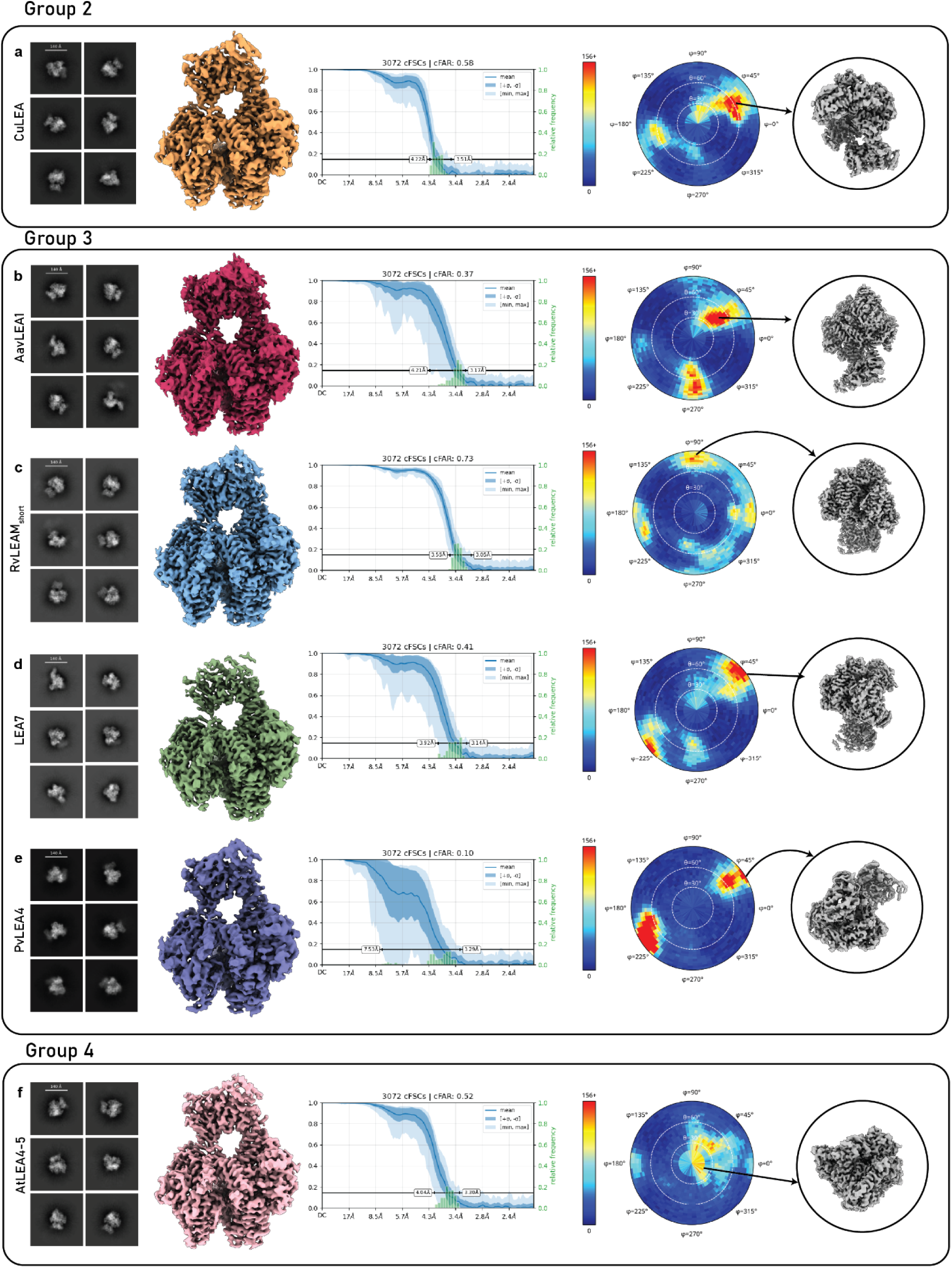
Each LEA protein induces a distinct orientation distribution for Polα-primase. Evaluation of each LEA protein with an apo-state Polα-primase. **(a)** Group 2 LEA protein, **(b-e)** group 3 LEA proteins, **(f)** group 4 LEA proteins. From left to right, the top 6 2-D classes, refined map, a cryoSPARC conical FSC, *cis*TEM orientation diagram, and an example preferred orientation mapped from the corresponding orientation diagram.

We first tested whether these LEA proteins could protect a fragile cryo-EM sample from AWI-induced damage. For this test, we used human DNA polymerase alpha-primase (Polα-primase), a multi-subunit protein complex that is severely damaged during cryo-EM grid preparation but can be rescued by AavLEA1 and RvLEAM_short_^15^. Polα-primase therefore provided a reference sample for asking whether the four new LEA proteins also confer AWI protection. We found that all four newly tested LEA proteins produced monodisperse Polα-primase particles of similar size (**Supplementary Figs. 8-12**), showing that LEA-mediated protection of this fragile complex is not limited to AavLEA1 and RvLEAM_short_.

Having established that all four newly tested LEA proteins protected Polα-primase from AWI-induced damage, we next evaluated their impact on the particle orientation distribution. The datasets were processed using the same workflow, conditions, and number of final particles. The four LEA proteins yielded apo-state Polα-primase maps with comparable global resolutions of 3.6 to 4.1 Å, as calculated by 3DFSC, similar to the resolutions previously obtained with AavLEA1 and RvLEAM_short_^15^ (**Fig. 3b-c**). Despite the similar map resolutions, the orientation distributions differed substantially across LEA proteins. AtLEA4-5 produced the most distinct distribution (**Fig. 3f**), whereas PvLEA4 had the most severe orientation bias (**Fig. 3e**). Isotropy metrics confirmed that all tested LEA proteins, except PvLEA4, produced sufficiently isotropic reconstructions (**Supplementary Table 3**). Thus, among the LEA proteins tested, AWI protection of Polα-primase was broadly shared, whereas the resulting particle-orientation distributions were LEA-dependent.

Together, these results suggest that AWI protection is a common property among the LEA proteins tested, while individual LEA proteins can offer distinct orientation distributions even for the same sample. This diversity in angular sampling provides the rationale for building a broader LEA protein catalog, not simply to identify more protective additives, but to expand the range of orientation distributions available for cryo-EM sample preparation.

### Post-collection merging is a more predictable strategy than LEA premixing for controlling particle-orientation distributions

The distinct Polα-primase orientation distributions produced by individual LEA proteins raised a practical question: could complementary LEA-induced distributions be combined to improve orientation coverage? Catalase provided an initial proof-of-concept that LEA proteins can alter particle orientations, but its preferred orientation was not severe enough to strongly compromise map isotropy or model building, and its symmetry can partially offset orientation gaps through symmetry-related views (**Fig. 1**). Polα-primase therefore provided a more sensitive test of LEA-dependent orientation diversity because it is an asymmetric C1 complex: each particle contributes only its observed orientation, making gaps in orientation coverage more consequential for map isotropy. We therefore compared two strategies: post-collection merging of particle stacks from individually collected LEA datasets and experimental premixing of multiple LEA proteins before plunge freezing.

We first assessed pairwise post-collection combinations of Polα-primase particle stacks by using equal number of particles from each dataset, repeating 3D refinement after combination, and quantifying the resulting orientation distributions using cFAR and SCF. We interpret these combinations as a test of orientation-distribution complementarity rather than as an attempt to improve model building, because individual LEA-containing datasets already yielded largely isotropic reconstructions. The extent of metric improvement depends on the relationship between the underlying orientation distributions. Datasets with similar distributions showed only modest gains upon combination, whereas more distinct but complementary pairings yielded larger improvements (**Supplementary Fig. 13**). The AavLEA1 plus AtLEA4-5 and CuLEA plus AtLEA4-5 combinations showed the greatest gains, consistent with AtLEA4-5 producing the most distinct distribution among six LEA proteins tested. In contrast, combinations that included RvLEAM_short_ showed modest or no improvement, reflecting the already favorable orientation distribution of the RvLEAM_short_ dataset alone.

After identifying complementary LEA-induced orientation distributions from pairwise combinations, we next combined all six LEA datasets to ask whether the full panel could further broaden orientation coverage and improve isotropic metrics. We merged equal number of particles from all six LEA datasets and repeated 3D refinement. As expected, the orientation distribution from the combined reconstructed map resembled a superposition of the individual distributions (**Fig. 4a, left**), however its isotropy metrics, cFAR of 0.69 and SCF of 0.877, were not substantially better than either individual or pairwise combined datasets (**Supplementary Tables 3 & 4)**. This suggests post-collection merging is most useful when the contributing distributions provide balanced and complementary orientation distributions.

**Figure 4:**
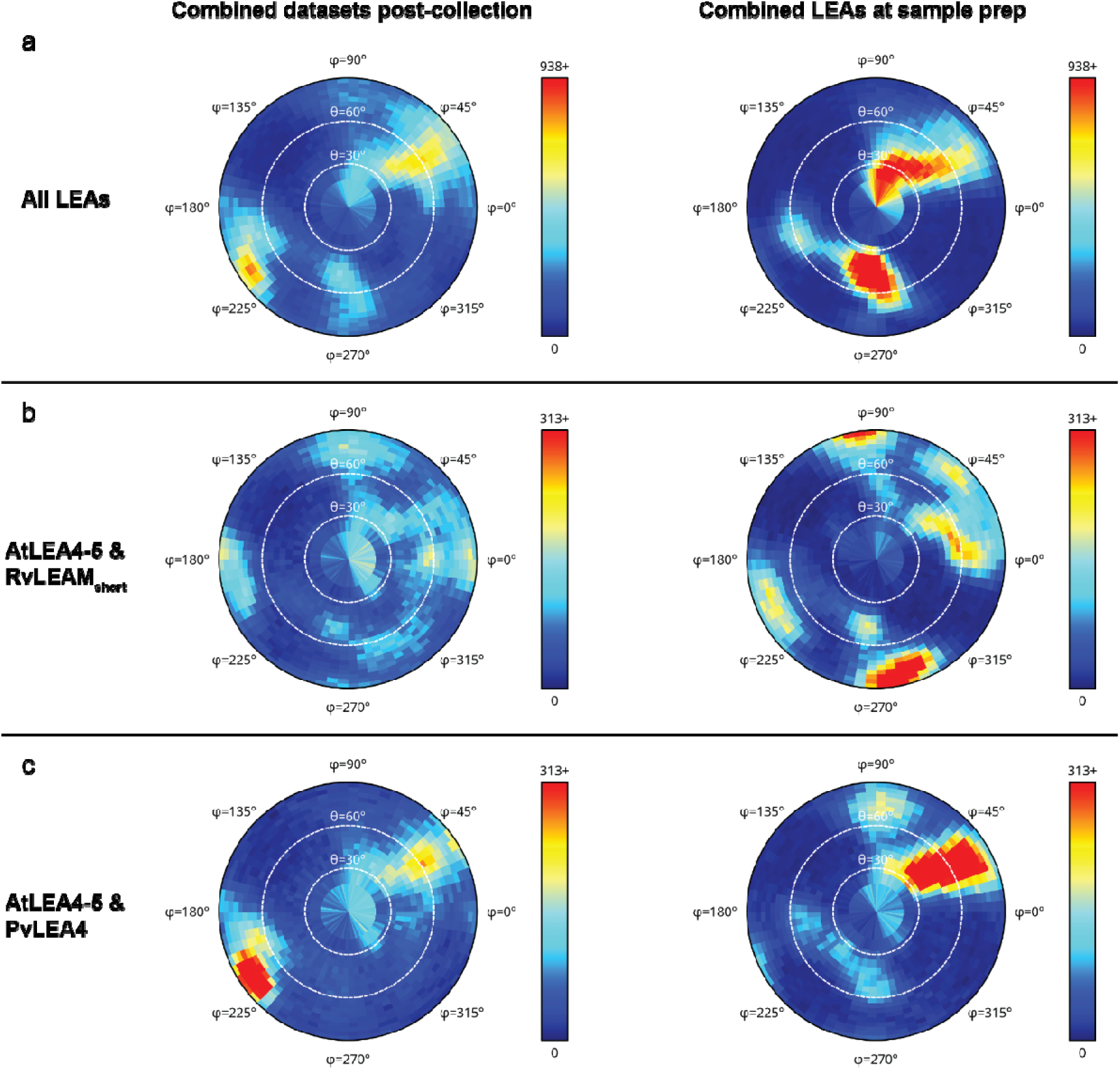
Comparison of premixed and post-collection LEA protein combinations on particle orientation. Orientation diagrams of each LEA protein combination with Polα-primase. **(a)** For all premixed (left) and post-collection (right) combination analyses, 270,000 particles were used to ensure equal comparison. **(b)** Premixed (left) and post-collection (right) AtLEA4-5 and RvLEAM_short_ combinations. 90,000 particles were used for each analysis. **(c)** Premixed (left) and post-collection (right) AtLEA4-5 and PvLEA4 combinations. 90,000 particles were used for each analysis.

We next asked whether post-collection merging could be reproduced experimentally by premixing LEA proteins during sample preparation. When all six LEA proteins were mixed with Polα–primase before plunge freezing at a total LEA concentration of 8 µM (1.33 µM each), the resulting orientation distribution was not a simple composite of the individual datasets (**Fig. 4a, right, Supplementary Fig. 14**). Instead, the premixed data was more biased that the all-six post-collection merge (cFAR = 0.43, SCF = 0.682), dominated by particle views resembling the AavLEA1, CuLEA, and PvLEA4 datasets (**Fig 3**), and contained a cluster of particle views not seen in the individual datasets. Pairwise LEA premixing showed the same non-additive behavior. For AtLEA4-5 plus RvLEAM_short_, post-collection merging improved orientation distribution (**Fig. 4b, left**; cFAR = 0.75, SCF = 0.935), as expected from their complementary orientation distributions. However, premixing produced a more biased distribution resembling RvLEAM_short_ alone (**Fig. 4b, right, Supplementary Fig. 15**; cFAR = 0.40, SCF = 0.700). For AtLEA4-5 plus PvLEA4, post-collection merging preserved the features of both individual distributions (**Fig. 4c, left**; cFAR = 0.51, SCF = 0.865), whereas premixing produced a different distribution with a new preferred orientation and a bias toward PvLEA4-like views (**Fig. 4c, right, Supplementary Fig. 16**; cFAR = 0.49, SCF = 0.725). Thus, post-collection merging predictably leverages complementary LEA-induced orientation distributions, whereas LEA premixing can produce non-additive effects, including dominance by one LEA condition or emergence of new preferred orientations.

## Discussion

Preferential orientation at the AWI is a major limitation in single-particle cryo-EM, preventing adequate angular sampling coverage and leading to suboptimal 3D reconstructions for many samples. Building on our previous work on LEA proteins protecting samples from AWI-induced damage^15^, we demonstrate here that they can also modulate particle orientation in a sample-dependent manner. This effect can be leveraged to improve orientation distributions and isotropy of reconstructed maps. This approach compares favorably with existing strategies for mitigating preferential orientation along several practical considerations. Compared with detergent-based approaches such as CHAPSO^11,14^ or the recently reported DM^41^, LEA protein application requires substantially lower sample concentrations, as little as 1 µM of target protein, compared to 5 to over 100 µM typically needed when detergents are used. This reduced concentration requirement becomes particularly advantageous for samples that are difficult to obtain or cannot be concentrated to sufficiently high levels for detergent use. In addition, LEA proteins are currently the only protein-based additives available, which are less likely to interact with target protein samples than small-molecule detergents^14^.

Selecting a LEA protein for a new cryo-EM target requires empirical screening because LEA effects on particle orientation are sample-specific, and individual LEA proteins can produce different orientation distributions. At present, no predictive framework exists to match a given target with the optimal LEA protein. We therefore recommend evaluating the six LEA proteins tested here independently during initial cryo-EM sample screening. LEA conditions that yield monodisperse particles of the expected size and shape can then be advanced to small-dataset collection, followed by 3D reconstruction using a standard SPA workflow. Orientation distributions from these datasets can be compared to identify the LEA protein that provides the most complete orientation coverage for the target. If no single LEA protein provides sufficient coverage, datasets from LEA protein conditions with complementary orientation distributions can be merged during data processing. Larger datasets can then be collected for the selected condition, or conditions, to obtain the highest-resolution reconstruction achievable for that target.

Looking ahead, future work should continue to expand the LEA protein catalog, establish empirical guidelines linking LEA protein properties to orientation outcomes, and evaluate whether LEA proteins can be combined with one another or with other AWI-mitigating strategies during grid preparation. These efforts would further establish LEA proteins as a versatile toolkit for mitigating preferred orientation during cryo-EM sample preparation.

## Methods

### Expression and purification of LEA proteins

Expression and purification of AavLEA1 and RvLEAM_short_ as described previously^15^. The sequences of CuLEA, LEA7, PvLEA4 and AtLEA4-5 were identified in the literature, and the genes were ordered from IDT, USA. The genes were inserted into a pET15b vector with a 6x-HIS tag and a GST-PreScission cut site. All recombinant plasmids were expressed in *Escherichia coli* BL21 (DE3) cells. A single bacterial colony containing the transformed plasmid was cultured overnight in 2 mL of Luria Bertani broth (LB) with 100 µg mL^−1^ carbenicillin at 37°C. The starter culture was then used to inoculate 1 L of Terrific Broth Auto-Induction-Medium (TB-AIM) (Boca Scientific, USA), supplemented with the same antibiotic. The culture was kept at 30°C for 16 hours shaking at 230 rpm before harvesting.

Cells were harvested by centrifugation, resuspended in lysis buffer (50 mM HEPES pH 7.5, 300 mM NaCl, 20 mM Imidazole, 1 mM DTT or TCEP, 1 mM PMSF) and lysed via sonication. Lysate was then incubated at 85°C for 30 minutes, followed by a 15-minute incubation on ice. Cell debris was then removed by centrifugation. The clarified lysate was then incubated with pre-equilibrated nickel-NTA resin (Qiagen, Germany) and stirred for 1 hour at 4°C. The protein bound resin was then washed 3 times with lysis buffer. Protein was then eluted with 10 mL of elution buffer (lysis buffer supplemented with 300 mM Imidazole) with a gravity flow column. The proteins were buffer exchanged and concentrated with SEC buffer (50 mM HEPES pH 7.5, 300 mM NaCl, 1 mM TCEP, 10% glycerol) and incubated overnight with GST-PreScission protease at 4°C to cleave off the HIS-tag.

The protein was then run on a Superdex 75 10/300 size-exclusion chromatography (SEC) column (Cytiva, USA) pre-equilibrated with the SEC buffer. Eluted fractions were analyzed by SDS-PAGE. Fractions were then chosen based on size and purity to be pooled, concentrated and snap-frozen with liquid nitrogen in 5 µl aliquots and stored at −80°C for future use. The protein concentrations of each aliquot were determined using the Beer-Lambert equation, with absorbance measurements obtained from a NanoDrop spectrophotometer (Thermo Fisher, USA) and extinction coefficients calculated based on protein sequence.

### Production of recombinant human DNA polymerase alpha-primase

Recombinant human Polα-primase was expressed and purified in *Trichoplusia ni (Tni)* cells (Expression Systems, USA) through baculovirus infection, as described previously^15,42^. Each of the four subunits was in its own baculovirus and co-expressed. After 66 hours of infection, the Tni cells were harvested and lysed. The cellular lysate was clarified and incubated with Ni-NTA agarose resin (Qiagen, Germany) to capture three of the His-tagged subunits. The elute was then incubated with Strep-Tactin XT 4Flow-resin (IBA LifeScience, Germany) to capture the strep-tagged subunit. The final purified Polα-primase was confirmed via SDS-PAGE analysis.

### Preparation of HA trimer and human erythrocyte Catalase

HA trimer (H3N2)(A/Hong Kong/1/1968) was purchased from My BioSource, USA (MBS434205). The protein was split into 5 µl aliquots, snap-frozen and stored at −20°C.

Human Erythrocyte Catalase was purchased from Sigma-Aldrich, USA (C3556). The lyophilized protein was resuspended in 25 mM HEPES pH 7.5, 150 mM NaCl, and 1 mM TCEP and snap-frozen in 5 µl aliquots. Protein was then stored at −20°C until use.

### Cryo-EM grid preparation and plunge freezing

All samples were thawed on ice just prior to grid preparation. C-flat R 1.2/1.3 300 mesh Cu holey carbon grids (CF313-50) were glow discharged using a PELCO EasiGlow (Ted Pella Inc, USA) at 15 mA for 30 s with a 10 s hold. Grids were used within 30 minutes of glow-discharging. Protein samples were diluted to the working concentrations and, when applicable, mixed with LEA proteins immediately before vitrification. All Catalase and Polα-primase datasets had a final sample protein concentration of 1 µM and 8 µM of LEA protein, where applicable. For datasets containing combinations of LEA proteins, each LEA protein was present at equal concentration with the total LEA concentration maintained at 8 µM. The HA trimer – AavLEA1 dataset contained 1.5 µM HA trimer and 12 µM AavLEA1. Finally, the HA trimer – RvLEAM_short_ dataset contained 1.5 µM HA trimer and 6 µM RvLEAM_short_. All samples were prepared in 25 mM HEPES pH 7.5, 150 mM NaCl and 1 mM TCEP. A 3.5 µl aliquot of the sample was applied to the glow-discharged grids, blotted for 2-3.5 seconds, and plunge-frozen into liquid ethane using either a GraVitri Plunge Cooler (MiTeGen, USA) or Vitrobot Mark IV (Thermo Fisher Scientific, USA). The GraVitri Plunge Cooler was used with an Ethane Cryostat^43^ with the temperature set at −180°C, blotting was done at room temperature and approximately 60% humidity. Grids prepared with the Vitrobot Mark IV were blotted at 4°C and 95% humidity.

### Cryo-EM data collection

Data collections and grid screenings were performed on a Talos Arctica 200 kV TEM (Thermo Fisher Scientific, USA) equipped with a Gatan BioQuantum K3 direct electron detector (Gatan, USA). EPU (Thermo Fisher Scientific, USA) was used for automatic data acquisition. All datasets were collected at a pixel size of 1.064 Å/pixel with a total dose of 50 e^−^Å^−2^ distributed over 40 frames. Data was recorded in CDS counting mode with a 20 eV energy filter slit inserted. The defocus range was set between −1 and −2.5 µm in 0.25-µm increments.

### Cryo-EM data processing

All datasets were processed using cryoSPARC^29^.

### 1 µM Catalase dataset

A total of 2,912 movies were collected. After micrograph curation, 654,360 particles were selected from 500 micrographs and extracted at 4× binning (4.3□Å/pixel). These particles underwent 2D classification and were used to generate picking templates. Template-based picking was then applied to the full movie stack, yielding 1,986,990 particles, which were extracted at 4× binning. Following two additional rounds of 2D classification, 182,472 particles remained and were re-extracted at the unbinned pixel size of 1.064□Å/pixel. Particles then underwent reference-based motion correction and global CTF refinement, followed by heterogeneous refinement into four classes. The best class, containing 112,544 particles, was subjected to non-uniform refinement with D2 symmetry, producing a reconstruction at 3.16□Å global resolution. A random subset of 90,000 particles was further refined with non-uniform refinement and D2 symmetry, yielding a global resolution of 3.06□Å. Half-maps were used to compute the 3DFSC, which reported a resolution of 3.24 Å.

### 1 µM Catalase, 8 µM AavLEA1 dataset

A total of 1,168 movies were collected. After micrograph curation, 399,852 particles were selected using templates generated in the 1 µM Catalase dataset and extracted at 4× binning (4.3□Å/pixel). These particles underwent two rounds of 2D classification, and the remaining 151,574 particles were re-extracted at the unbinned pixel size of 1.064 Å/pixel. Particles then underwent reference-based motion correction and were classified into four classes via heterogeneous refinement. The best two classes, comprising of 89,819 particles, were subjected to global CTF refinement followed by non-uniform refinement with D2 symmetry yielding a 2.8 Å map reconstruction. Half-maps were used to compute the 3DFSC, which also reported a resolution of 2.8 Å.

### 1 µM Catalase, 8 µM RvLEAM_short_ dataset

A total of 3,625 movies were collected. After micrograph curation, 1,163,787 particles were selected using templates generated in the 1 µM Catalase dataset and extracted at 4× binning (4.3□Å/pixel). These particles underwent two rounds of 2D classification, and the remaining 340,332 particles were re-extracted at the unbinned pixel size of 1.064 Å/pixel. Particles were then split into four classes via heterogeneous refinement, and 241,262 particles from the top three classes underwent another round of 2D classification. These particles were then subjected to reference-based motion correction, global CTF refinement and another round of heterogeneous refinement with 4 classes. The top class was chosen which consisted of 131,430 particles and underwent non-uniform refinement with D2 symmetry resulting in a 3.2 Å map. A subset of 90,000 randomly chosen particles underwent non-uniform refinement resulting in a resolution of 3.3 Å. Half-maps were used to compute the 3DFSC resolution which also reported a 3.3 Å map.

### 1 µM HA trimer dataset

A total of 4,499 movies were collected. After micrograph curation, 1,340,513 particles were selected using the cryo-SPARC deep inference job (trained on the 1.5 µM HA trimer, 12 µM AavLEA1 dataset) and extracted at 4x binning (4.3 Å/pixel). These particles underwent two rounds of 2D classification, and the remaining 403,088 particles were re-extracted at 1.064 Å/pixel. These particles then underwent reference-based motion correction, global CTF refinement followed by non-uniform refinement using C3 symmetry. This yielded a map with 3.2 Å global resolution. A subset of 87,122 particles were randomly chosen from this dataset for the final reconstruction, giving a 3.4 Å map. Half-maps were used to compute the 3DFSC resolution which reported a 3.5 Å map.

### 1.5 µM HA trimer, 12 µM AavLEA1 dataset

A total of 8,965 movies were collected. After micrograph curation, 5,946,736 particles chosen using template picker and were extracted at 4x binned. These particles underwent four rounds of 2D classification and a round of ab-initio reconstruction before they were re-extracted at 1.064 Å/pixel. The remaining 587,981 particles underwent heterogeneous refinement where they were split into four classes. Classes 1, 2 and 3 were chosen, bringing the particles to 405,557. These particles then underwent a second round of heterogeneous refinement and were split into four classes again. The 98,685 particles in class 3 were chosen and underwent another round of 2D classification to clean up the particle stack. The remaining particles were used to train the Deep Train machine learning model in cryo-SPARC. All micrographs were de-noised, and particles were re-selected from the de-noised micrographs using Deep Inference. The 1,725,541 re-selected particles were extracted at 4x binned and underwent two rounds of 2D classification and then re-extracted at 1.064 Å/pixel. These particles were then split into four classes via heterogeneous refinement. Only particles from class 3 of this job were selected and underwent reference-based motion correction, global CTF refinement and finally non-uniform refinement with C3 symmetry. The final map had a global resolution of 3.7 Å and half maps were used to compute the 3DFSC resolution which also reported a 3.7 Å map.

### 1.5 µM HA trimer, 6 µM RvLEAM_short_ dataset

A total of 6,136 movies were collected. Following micrograph curation and de-noising, the deep inference job was run using particles from the 1.5 µM HA trimer, 12 µM AavLEA1 dataset as training. The 1,545,514 particles were extracted at 4x binning and underwent two rounds of 2D classification. The remaining 294,832 particles were then extracted at 1x binning (1.064 Å/pixel) and were split into four classes using heterogeneous refinement. Classes 1 and 2 were selected yielding 172,523 particles. These particles underwent reference motion correction, global CTF refinement and non-uniform refinement with C3 symmetry. This produced a map with a global resolution of 3.4 Å. A random subset of 87,122 particles were selected from this particle stack and underwent a final round of non-uniform refinement using C3 symmetry. This resulted in a 3.6 Å map. The 3DFSC resolution, calculated using half maps, also resulted in a 3.6 Å final map.

### 1 µM Polα-primase, 8 µM RvLEAM_short_ dataset

A total of 5,211 movies were collected. Following micrograph curation, particles from 500 micrographs were selected using blob-picker and extracted at 4x binning. The 155,182 particles underwent 2D classification and then were used as templates for template picking for the rest of the micrographs. 2,865,601 particles were chosen from the remaining micrographs and were extracted at 4x binning. All particles underwent 2D classification and the remaining 970,403 particles were then subjected to ab-initio reconstruction with 4 classes. Particles from class 0 were chosen and extracted at 1.064 Å/pixel, underwent another round of 2D classification followed by reference-based motion correction. The remaining 498,863 particles were split into two more classes through ab-initio reconstruction. Class 1 was chosen and was further split into four more classes via heterogeneous refinement. From here, particles from class 1 were chosen, and underwent global CTF refinement followed by non-uniform refinement. This stack of 357,409 particles produced a 3.0 Å global resolution map. A stack of 45,000 particles was randomly chosen from this stack and underwent a final round of non-uniform refinement with C1 symmetry. This gave a final global resolution of 3.4 Å, and a 3DFSC resolution of 3.5 Å calculated from half-maps.

### 1 µM Polα-primase, 8 µM AtLEA4-5 dataset

A total of 3,934 movies were collected. Following micrograph curation, 1,216,282 particles were selected from template picking and extracted at 4x binning. These particles underwent 2D classification, cutting the particle stack down to 479,762 particles. These particles were split into four classes via ab-initio reconstruction. Particles from class 1 were extracted at 1.064 Å and underwent another round of 2D classification. The remaining 231,541 particles underwent a second round of ab-initio reconstruction with four classes. Classes 2 and 3 were selected and the 188,027 particles underwent reference-based motion correction, global CTF refinement and non-uniform refinement with C1 symmetry, resulting in a 3.4 Å global resolution map. A random subset of 45,000 particles were selected from this particle stack and underwent a final round of non-uniform refinement resulting in a 3.4 Å global resolution map and a 3.83 Å 3DFSC resolution, calculated from half-maps.

### 1 µM Polα-primase, 8 µM CuLEA dataset

A total of 4,551 movies were collected. Following micrograph curation, 2,055,454 particles were selected via template picking and were extracted at 4x binning. Particles then underwent two rounds of 2D classification, and the remaining 322,513 particles were split into four classes in ab-initio classification. Particles from class 0 were extracted at 1.064 Å and underwent another round of 2D classification. The remaining 232,531 particles were split into two classes through ab-initio reconstruction and particles from class 1 were sent through a round of heterogeneous refinement with four classes. Particles from class 2 and 3 were selected and underwent reference-based motion correction, global CTF refinement and non-uniform refinement with C1 symmetry. These 164,921 particles resulted in a 3.7 Å map. A random subset of 45,000 particles were selected from this particle stack and underwent a final round of non-uniform refinement with C1 symmetry resulting in a 4.0 Å global resolution map, and a 4.1 Å 3DFSC map, calculated from half-maps.

### 1 µM Polα-primase, 8 µM LEA7 dataset

Due to some microscope issues, this dataset was collected on two separate days, one of which the micrographs were gain-normalized and on the second day the micrographs were not. Because of this, each sub-set of this dataset was processed separately until the final refinement steps.

A total of 1,703 non-gain normalized movies were collected, underwent micrograph curation and 920,141 particles were picked via blob picking and extracted at 4x binning. These particles underwent 2D classification followed by ab-initio reconstruction where they were split into 4 classes. Classes 1 and 2 were selected and the 250,687 particles from these classes were extracted with recentering at 1.064 Å/pixel and underwent reference motion correction and global CTF refinement. These particles then were refined in a non-uniform refinement job giving a global resolution of 3.4 Å.

2,116 gain-normalized movies were collected, underwent micrograph curation and template picking (using templates from the non-gain normalized dataset), and extracted at 4x binning. The 1,618,851 extracted particles underwent a round of 2D classification, and the remaining 575,185 particles were subjected to ab-initio reconstruction with 4 classes. Particles from class 3 were selected and extracted with recentering at 1.064 Å/pixel followed by reference-based motion correction and global CTF refinement. The final 397,165 particles produced a 3.2 Å map from non-uniform refinement.

The particles from both gain-normalized and non-gain normalized subsets were then pooled yielding 647,622 particles. This particle stack was split into four classes from a 3D classification job. Classes 0, 2 and 3 were selected and a non-uniform refinement job resulted in a 3.1 Å global resolution with 567,915 particles. A subset of 45,000 particles were randomly selected from this particle stack and underwent a final round of non-uniform refinement. This resulted in a 3.5 Å global resolution map and a 3DFSC resolution of 3.6 Å calculated using half maps.

### 1 µM Polα-primase, 8 µM PvLEA4 dataset

A total of 2,876 movies were collected. Following micrograph curation, 1,310,646 particles were selected through template picking, and were extracted at 4x binning. These particles underwent two rounds of 2D classification followed by ab-initio reconstruction with 4 classes. Particles from classes 1 and 2 were extracted at 1.064 Å/pixel with recentering. The 70,018 extracted particles underwent a second round of ab-initio reconstruction where they were split into two classes. Particles from class 1 were selected and underwent reference-based motion correction, global CTF refinement and finally, non-uniform refinement with C1 symmetry. The remaining 45,009 particles produced a 4.1 Å global resolution final map, which was also reported by the 3DFSC resolution using half-maps.

### 1 µM Polα-primase, 8 µM all LEAs combined dataset

A total of 4,087 movies were collected. Following micrograph curation, 2,707,076 particles were selected through blob picking and extracted at 4x binning. These particles underwent three rounds of 2D classification followed by ab-initio reconstruction with 4 classes. Particles from classes 2 and 3 underwent another round of 2D classification and were then extracted at 1.064 Å/pixel with recentering. The 567,164 particles then underwent heterogeneous refinement with 4 classes and the 371,796 particles in class 2 underwent three more round of 2D classification, reference-based motion correction and global CTF refinement. The remaining 325,777 particles underwent non-uniform refinement with C1 symmetry resulting 3.7 Å map. From this particle stack, a random 270,000 particles were chosen for equal comparison to the computational combined dataset and underwent a final round of non-uniform refinement resulting in a final global resolution of 3.7 Å.

### 1 µM Polα-primase, 8 µM RvLEAM_short_ and AtLEA4-5 dataset

A total of 6,120 movies were collected. Following micrograph curation, 8,222,213 particles were selected through template picker (using templates generated from the all-LEA dataset) and extracted at 5x binning. These particles underwent two rounds of 2D classification and a random subset of 1,000,000 particles from the remaining 2,630,576 particles were sent into ab-initio reconstruction with 4 classes. Each volume generated from the ab-initio reconstruction was used as the initial volumes to seed a heterogeneous refinement with all 2.6 million particles. The 1,645,482 particles from class 1 were extracted with recentering and underwent another round of heterogeneous refinement with 4 more classes. Particles from class 1 were selected again and underwent a final round of heterogeneous refinement with 4 classes again. Particles from classes 0, 2 and 3 from this job were selected and sent into an ab initio job with 2 classes for one more round of cleanup. The 942,990 particles in class 0 were selected and underwent reference-based motion correction, global CTF refinement and non-uniform refinement with C1 symmetry resulting in a 2.8 Å map. A randomized subset of 90,000 particles was selected from this stack and underwent a final round of non-uniform refinement resulting in a final global resolution of 3.2 Å.

### 1 µM Polα-primase, 8 µM PvLEA4 and AtLEA4-5 dataset

A total of 6,045 movies were collected. Following micrograph curation, 7,041,237 particles were selected through template picker (using templates generated from the all-LEA dataset) and extracted at 5x binning. The particles underwent two rounds of 2D classification, yielding 477,430 particles. This stack was then subjected to ab-initio reconstruction with 4 classes. Particles from class 2 were extracted at 1x binning with recentering and underwent a second round of ab-initio with 4 more classes. Particles from classes 0, 2 and 3 were selected for a round of 2D classification resulting in 230,209 particles. These particles underwent a final round of ab-initio reconstruction with 4 classes and classes 0-2 were selected. The remaining 175,093 particles underwent non-uniform refinement with C1 symmetry yielding a 3.7 Å map. A randomized subset of 90,000 particles was selected from this stack and underwent a final round of non-uniform refinement resulting in a final global resolution of 3.8 Å.

### Orientation distribution analysis

Orientation metrics were calculated using CryoSPARC orientation diagnostics cFAR and SCF^30^. We included 3DFSC resolution and sphericity^8^ in our analysis as well. Orientation diagrams for each dataset were generated with *cis*TEM^44^.

## Data availability

The described cryo-EM maps, models, and motion-corrected movies have been deposited in the Electron Microscopy Databank (EMDB), the Protein Data Bank (PDB), and the Electron Microscopy Public Image Archive (EMPIAR) under the following accession codes: Catalase control EMD-77717, PDB-360G, EMPIAR-XXXXX. Catalase-AavLEA1 EMD-77718, PDB-36OH, EMPIAR-XXXXX. Catalase-RvLEAM_short_ EMD-77719, EMPIAR-XXXXX. HA-trimer control EMD-77215, PDB-35UY, EMPIAR-XXXXX. HA-Trimer-AavLEA1 EMD-77216, PDB-35UZ, EMPIAR-XXXXX. HA-Trimer-RvLEAM_short_ EMD-77217, PDB-35VA, EMPIAR-XXXXX. Polα-primase-RvLEAM_short_ EMD-77627, EMPIAR-XXXXX. Polα-primase-AtLEA4-5 EMD-77625, EMPIAR-XXXXX. Polα-primase-CuLEA EMD-77626, EMPIAR-XXXXX. Polα-primase-PvLEA4 EMD-77629, EMPIAR-XXXXX. Polα-primase-LEA7 EMD-77628, EMPIAR-XXXXX. Polα-primase-all LEAs EMD-XXXXX, EMPIAR-XXXXX. Polα-primase-AtLEA4-5 & RvLEAM_short_ EMD-XXXXX, EMPIAR-XXXXX. Polα-primase-AtLEA4-5 & PvLEA4 EMD-XXXXX, EMPIAR-XXXXX.

## Author contributions

C.L. and K.M.A conceived the study. C.L. designed the experiments with support from K.M.A. K.M.A. made the recombinant proteins and cryo-EM grids and performed data collection. K.M.A. and C.L. analyzed the data, wrote the manuscript, and prepared the figures. T.G. supported manuscript writing, experimental designs, analysis, and figure preparation.

## Acknowledgements

The data for this work was collected at the Cryo-EM Research Center (CEMRC) in the Department of Biochemistry at UW-Madison. Thank you to the staff at the CEMRC for their support with the cryo-EM data collections. The Lim Lab is a member of the SBGrid consortium (www.sbgrid.org) and some analysis was performed using software compiled by SBGrid. Support for this research was provided to C.L. by the National Institutes of Health (NIH) (R01GM153806 and DP2GM150023) and through the Wisconsin Alumni Research Foundation (WARF) Draper Technology Innovation Fund (TIF). K.M.A. is supported by a NIH T32 pre-doctoral fellowship (T32GM130550) and the Denis R. A. and Martha Washburn Wharton Department Fellowship. T.G. is an Investigator of The Morgridge Institute for Research.

