## Supplementary Information for "Diversifying particle-orientation distributions in cryo-EM with LEA protein additives"

**Table of Contents**

**Supplementary Figure 1.** Data processing pipeline for catalase control dataset.

**Supplementary Figure 2.** Data processing pipeline for catalase and AavLEA1 dataset.

**Supplementary Figure 3.** Data processing pipeline for catalase and RvLEAM_short_ dataset.

**Supplementary Figure 4.** Combined catalase control and catalase with AavLEA1 post processing.

**Supplementary Figure 5.** Data processing pipeline for HA trimer control dataset.

**Supplementary Figure 6.** Data processing pipeline for HA trimer and AavLEA1 dataset.

**Supplementary Figure 7.** Data processing pipeline for HA trimer and RvLEAM_short_ dataset.

**Supplementary Figure 8.** Data processing pipeline for Polymerase α-primase and CuLEA dataset.

**Supplementary Figure 9.** Data processing pipeline for Polymerase α-primase and RvLEAM_short_ dataset.

**Supplementary Figure 10.** Data processing pipeline for Polymerase α-primase and LEA7 dataset.

**Supplementary Figure 11.** Data processing pipeline for Polymerase α-primase and PvLEA4 dataset.

**Supplementary Figure 12.** Data processing pipeline for Polymerase α-primase and AtLEA4-5 dataset.

**Supplementary Figure 13.** Polymerase α-primase post-processing combined dataset orientation metric heatmap.

**Supplementary Figure 14.** Data processing pipeline for Polymerase α-primase and all LEAs combined dataset.

**Supplementary Figure 15.** Data processing pipeline for Polymerase α-primase and AtLEA4-5 + RvLEAM_short_.

**Supplementary Figure 16.** Data processing pipeline for Polymerase α-primase and AtLEA4-5 + PvLEA4.

**Supplementary Table 1.** Orientation metrics for catalase datasets.

**Supplementary Table 2.** Orientation metrics for HA trimer datasets.

**Supplementary Table 3.** Orientation metrics for Polymerase α-primase datasets.

**Supplementary Table 4:** Orientation metrics for combined LEA- Polα-primase datasets.

**Supplementary Table 5.** Cryo-EM data collection, refinement and validation statistics for catalase.

**Supplementary Table 6.** Cryo-EM data collection, refinement and validation statistics for HA trimer.

**Supplementary Table 7.** Cryo-EM data collection, refinement and validation statistics for Polymerase α-primase.

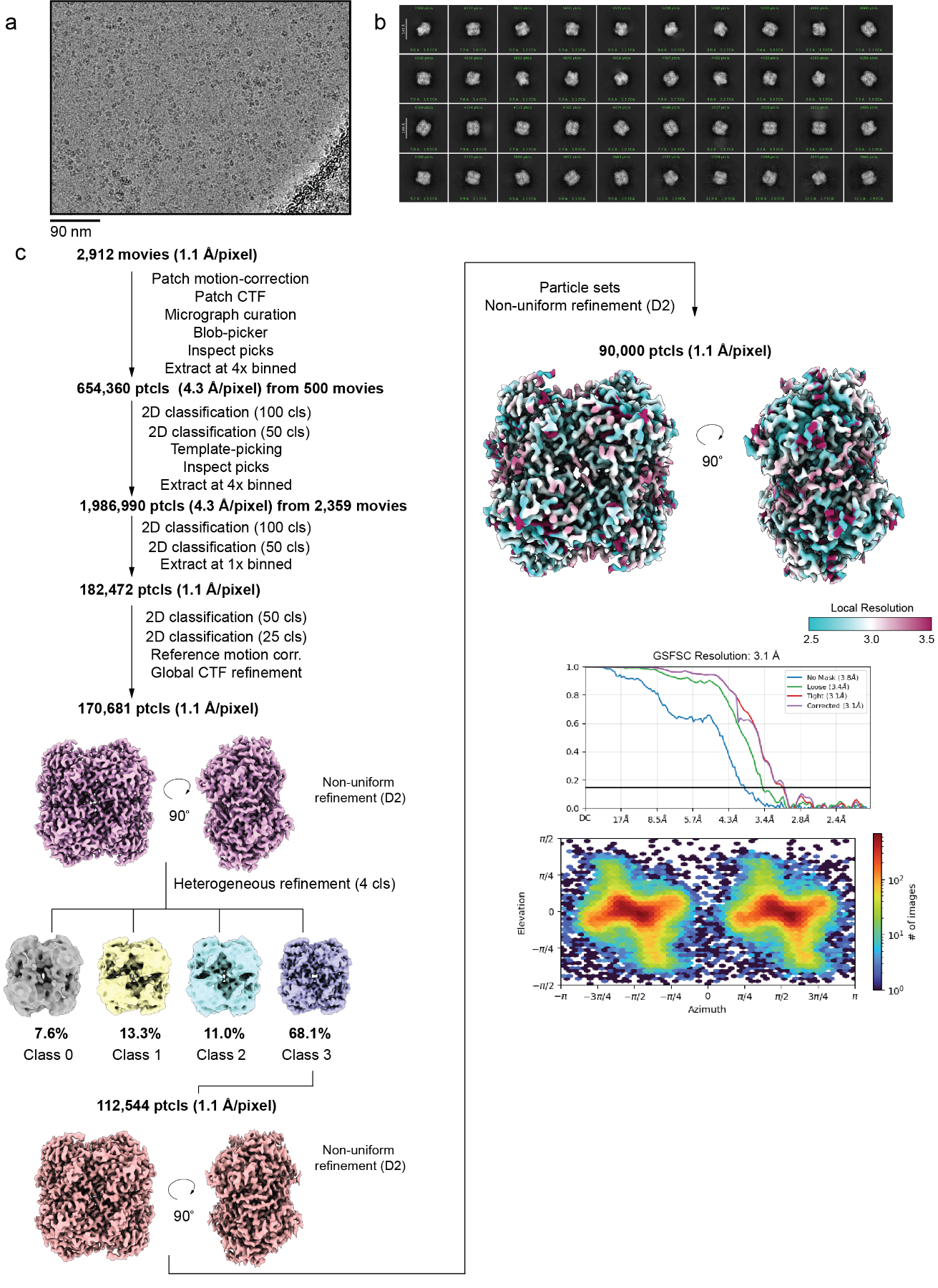

**Supplementary Figure 1** – **Data processing pipeline for catalase control dataset.** 1 µM catalase. **(a)** Representative micrograph. **(b)** Top 40 2D classes. **(c)** Cryo-EM processing pipeline used to obtain final map for catalase.

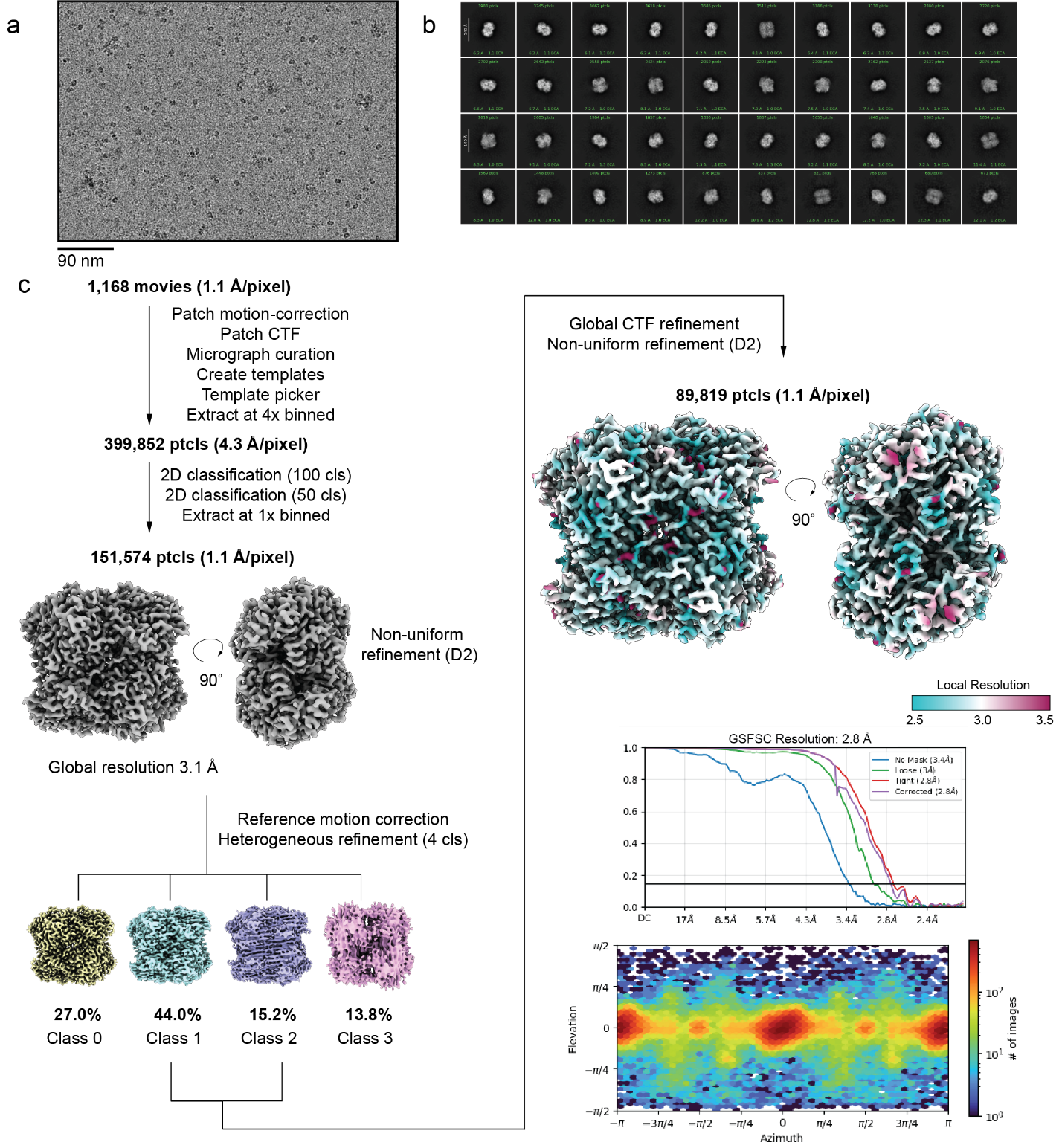

**Supplementary Figure 2** – **Data processing pipeline for catalase and AavLEA1 dataset.** 1 µM catalase with 8 µM AavLEA1. **(a)** Representative micrograph. **(b)** Top 40 2D classes. **(c)** Cryo-EM processing pipeline used to obtain final map for catalase.

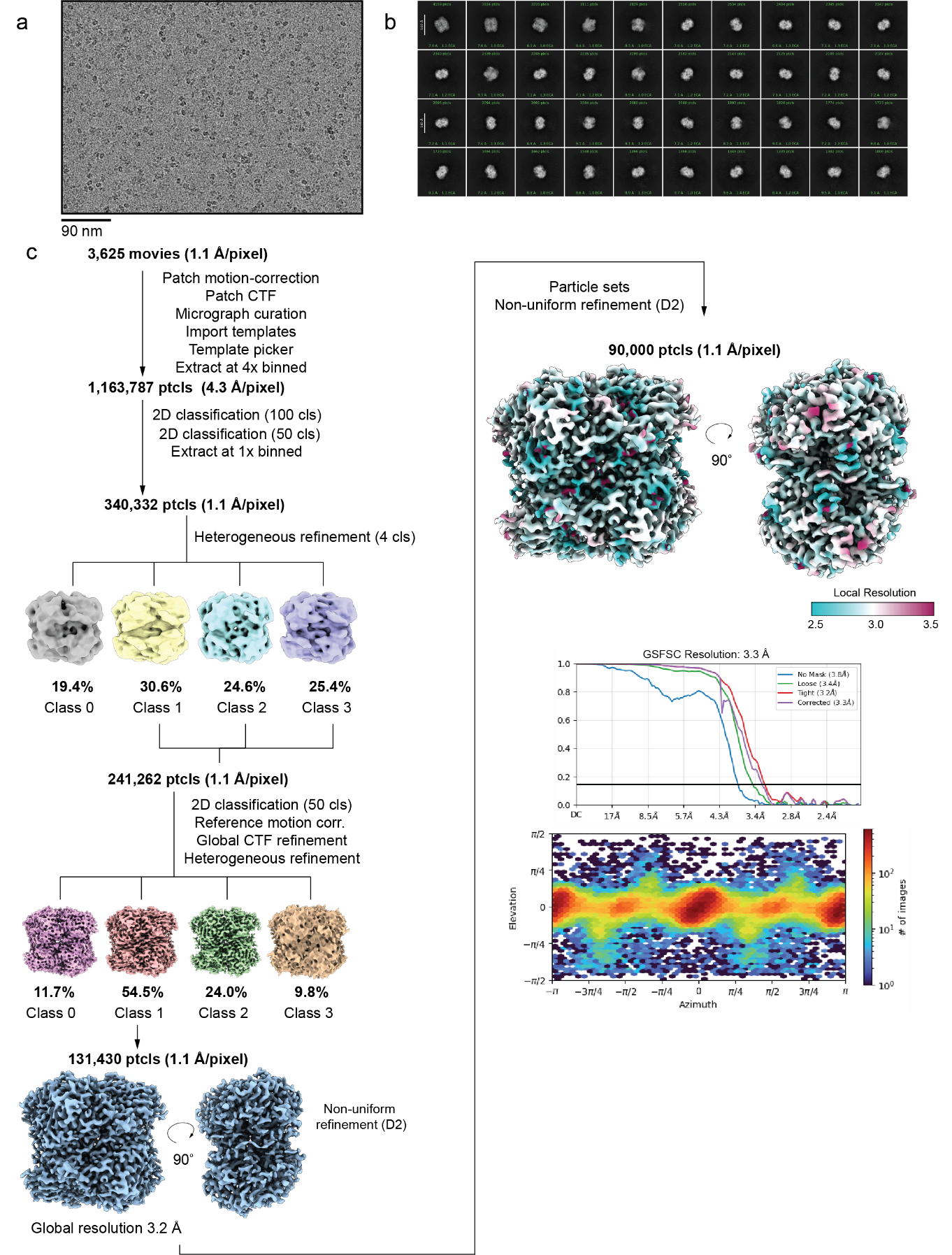

**Supplementary Figure 3** – **Data processing pipeline for catalase and RvLEAM_short_ dataset.** 1 µM catalase with 8 µM RvLEAM_short_. **(a)** Representative micrograph. **(b)** Top 40 2D classes. **(c)** Cryo-EM processing pipeline used to obtain final map for catalase.

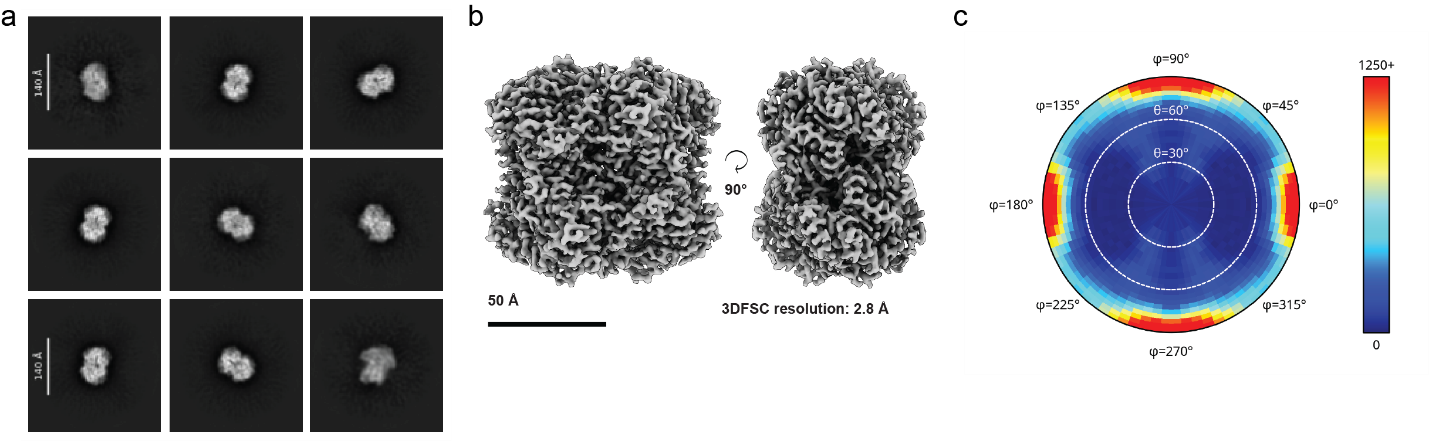
**Supplementary Figure 4 - Combined catalase control and catalase with AavLEA1 post processing. (a)** Representative 2D classes. **(b)** Final map. **(c)** Orientation diagram of combined dataset.

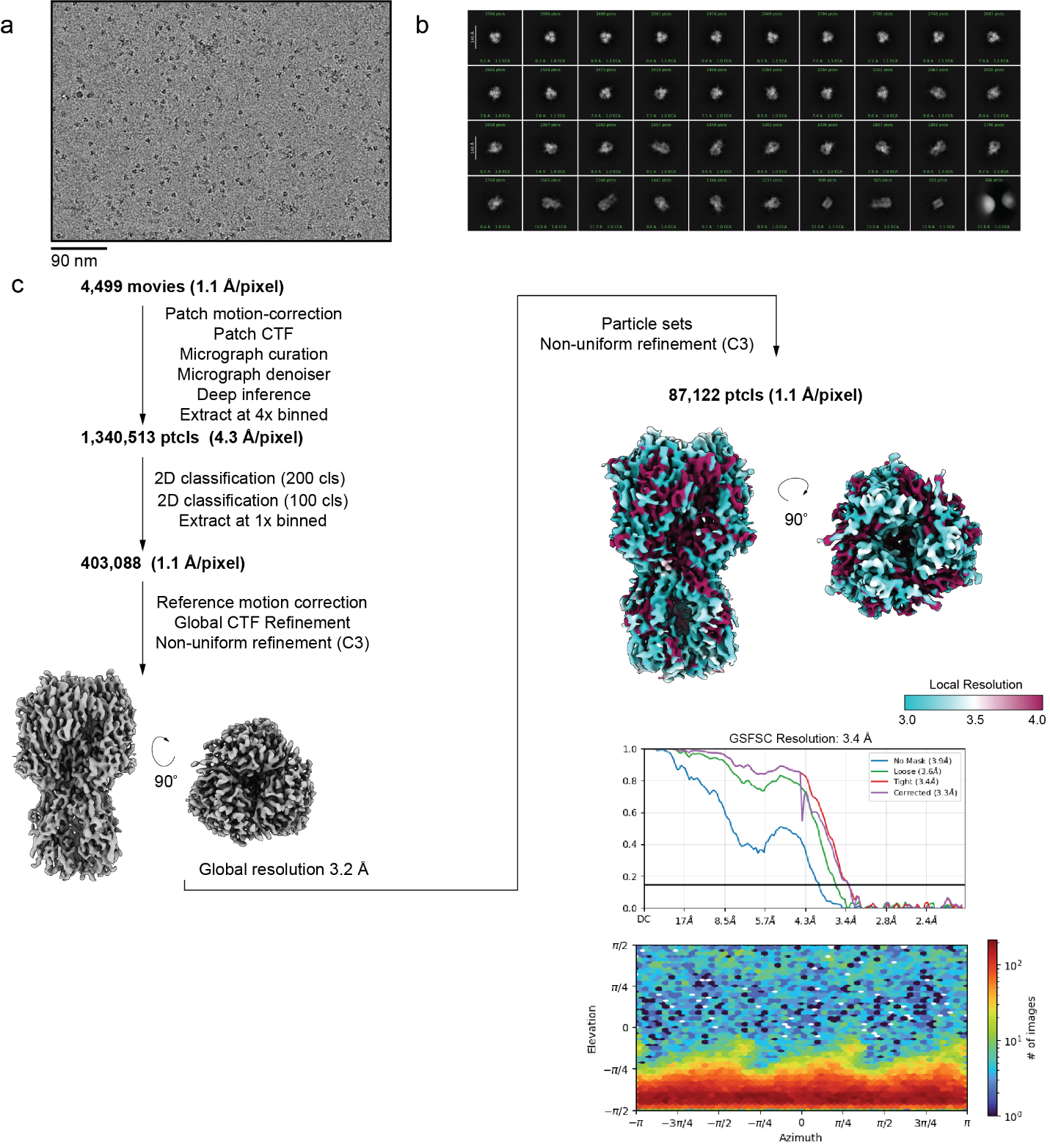

**Supplementary Figure 5 - Data processing pipeline for HA trimer control dataset.** 1 µM HA-trimer. **(a)** Representative micrograph. **(b)** Top 40 2D classes. **(c)** Cryo-EM processing pipeline used to obtain final map of HA-trimer.

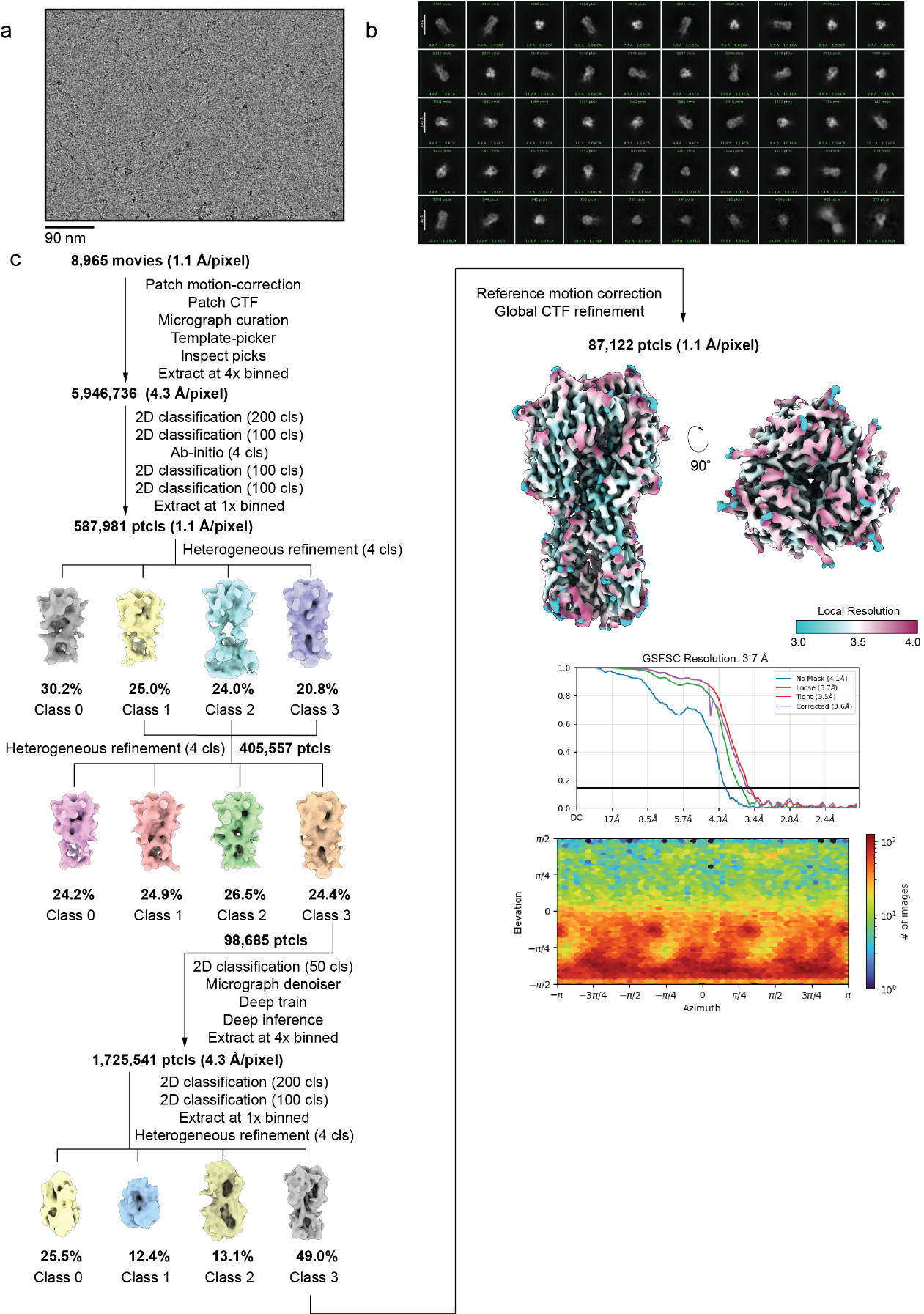

**Supplementary Figure 6 - Data processing pipeline for HA trimer and AavLEA1 dataset.** 1.5 µM HA-trimer with 12 µM AavLEA1. **(a)** Representative micrograph. **(b)** Top 40 2D classes. **(c)** Cryo-EM processing pipeline used to obtain final map of HA-trimer.

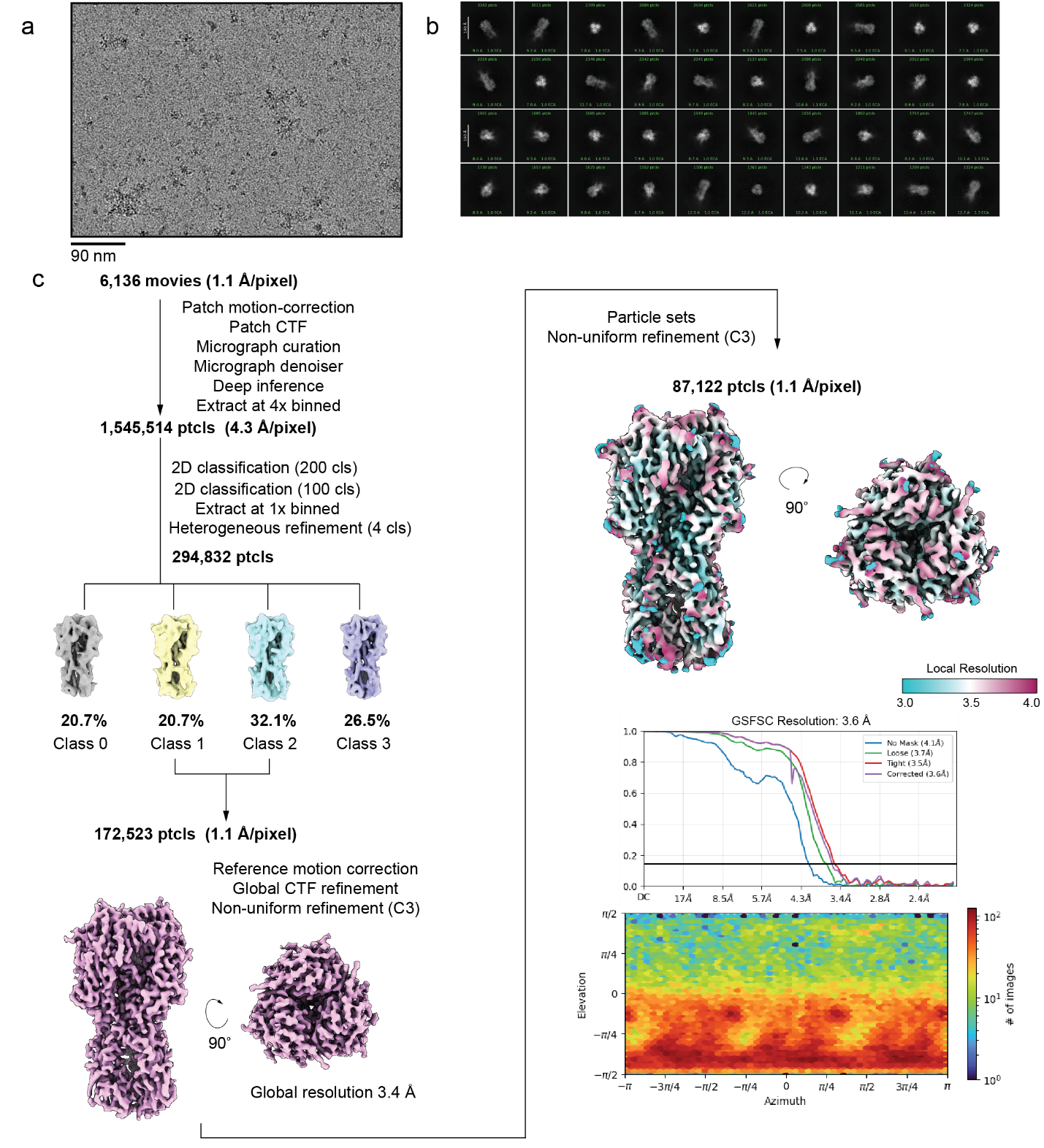

**Supplementary Figure 7 - Data processing pipeline for HA trimer and RvLEAM_short_ dataset.** 1.5 µM HA-trimer with 6 µM RvLEAM_short_. **(a)** Representative micrograph. **(b)** Top 40 2D classes. **(c)** Cryo-EM processing pipeline used to obtain final map of HA-trimer.

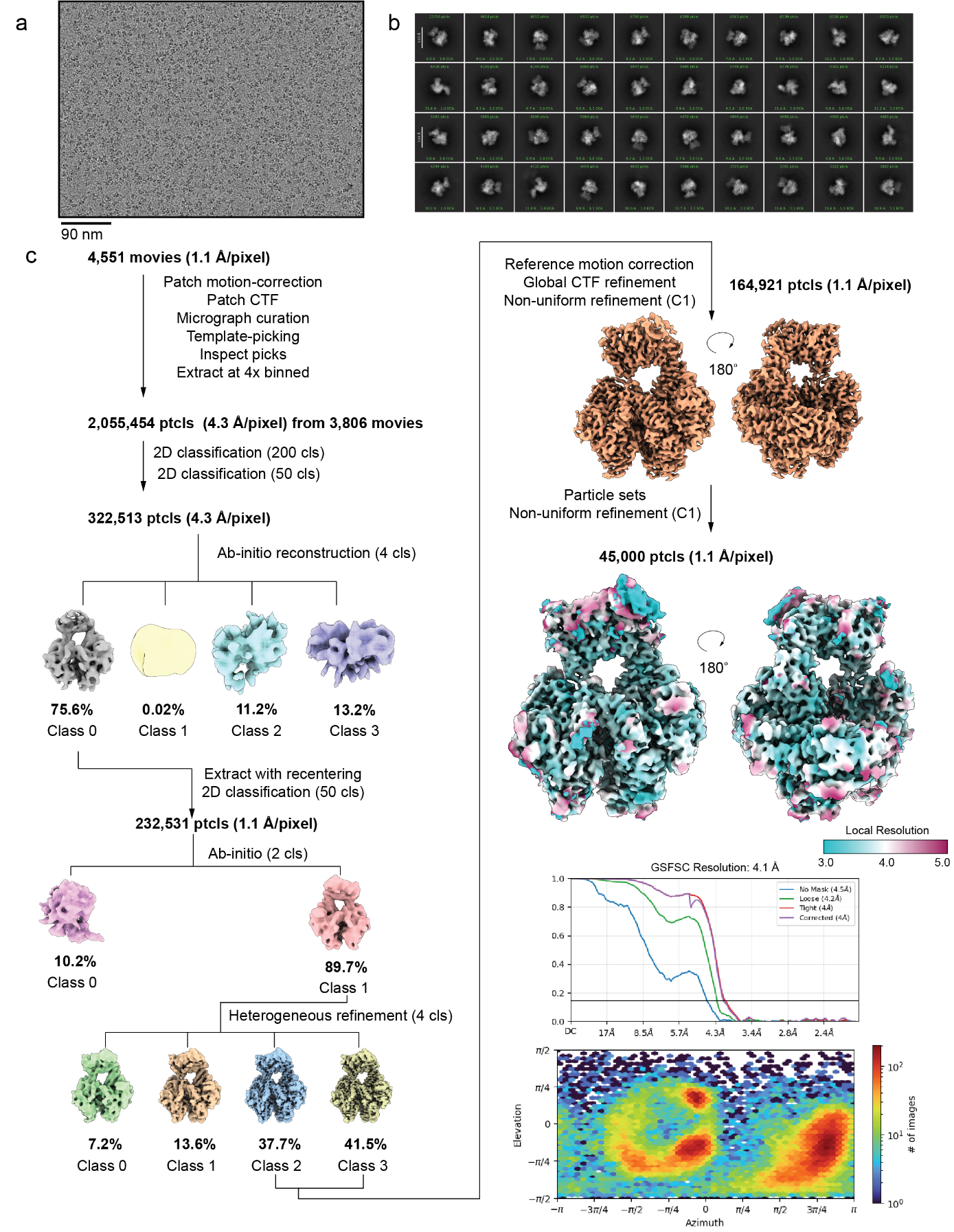
**Supplementary Figure 8 - Data processing pipeline for Polymerase α-primase and CuLEA dataset.** 1 µM Polα-primase with 8 µM CuLEA. **(a)** Representative micrograph. **(b)** Top 40 2D classes. **(c)** Cryo-EM processing pipeline used to obtain final map of Polα-primase.

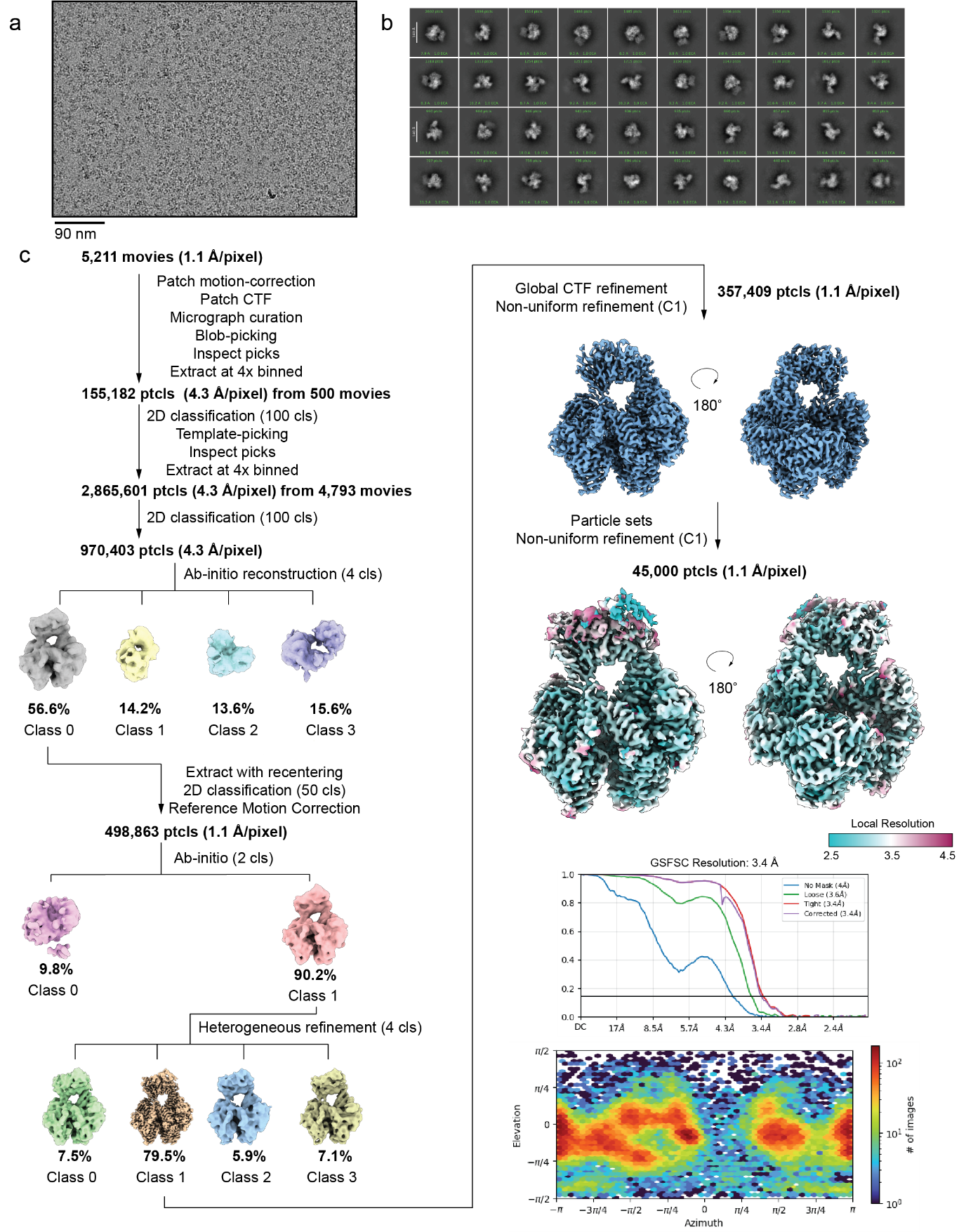

**Supplementary Figure 9 - Data processing pipeline for Polymerase α-primase and RvLEAM_short_ dataset.** 1 µM Polα-primase with 8 µM RvLEAM_short_. **(a)** Representative micrograph. **(b)** Top 40 2D classes. **(c)** Cryo-EM processing pipeline used to obtain final map of Polα-primase.

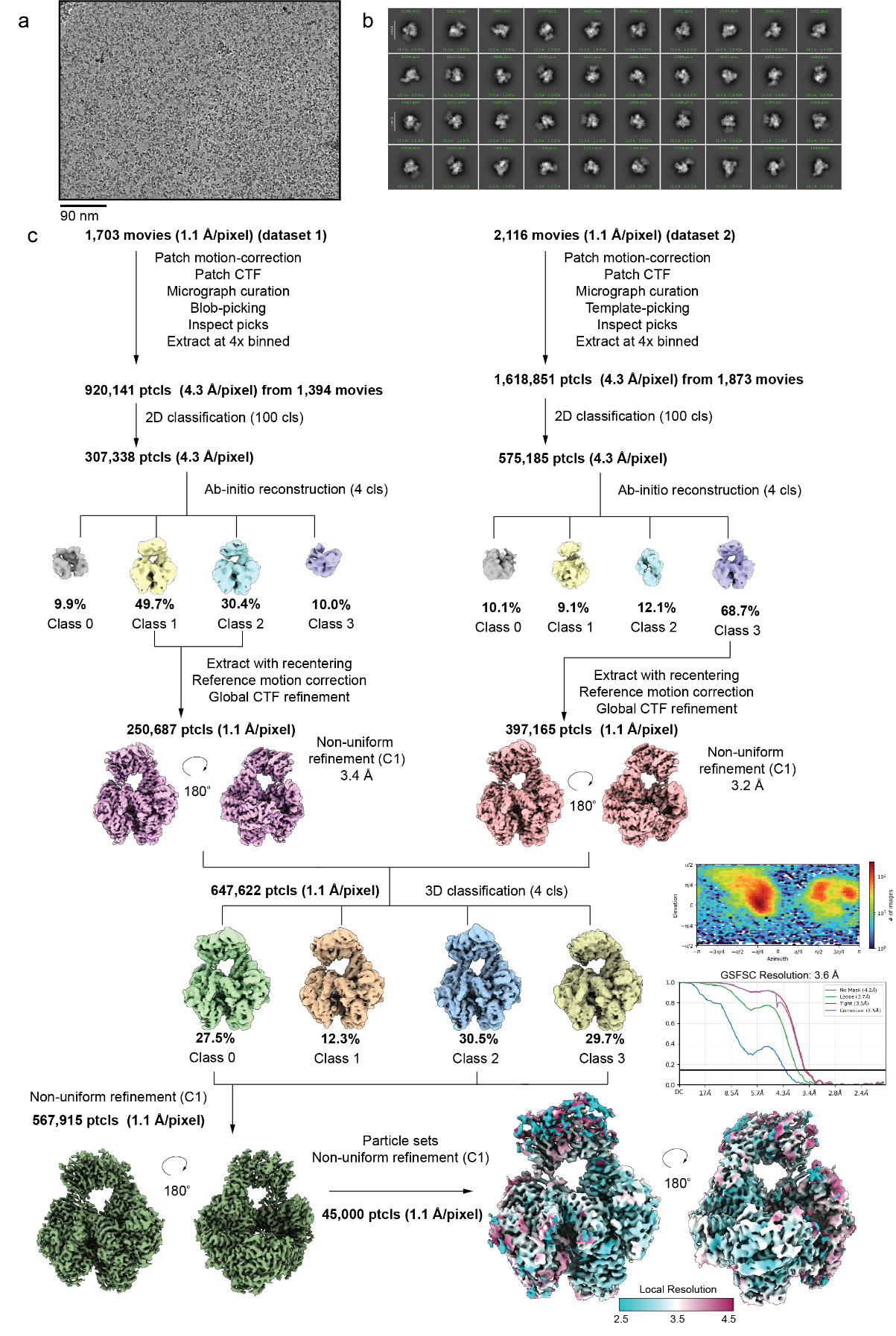

**Supplementary Figure 10 - Data processing pipeline for Polymerase α-primase and LEA7 dataset.** 1 µM Polα-primase with 8 µM LEA7. **(a)** Representative micrograph. **(b)** Top 40 2D classes. **(c)** Cryo-EM processing pipeline used to obtain final map of Polα-primase.

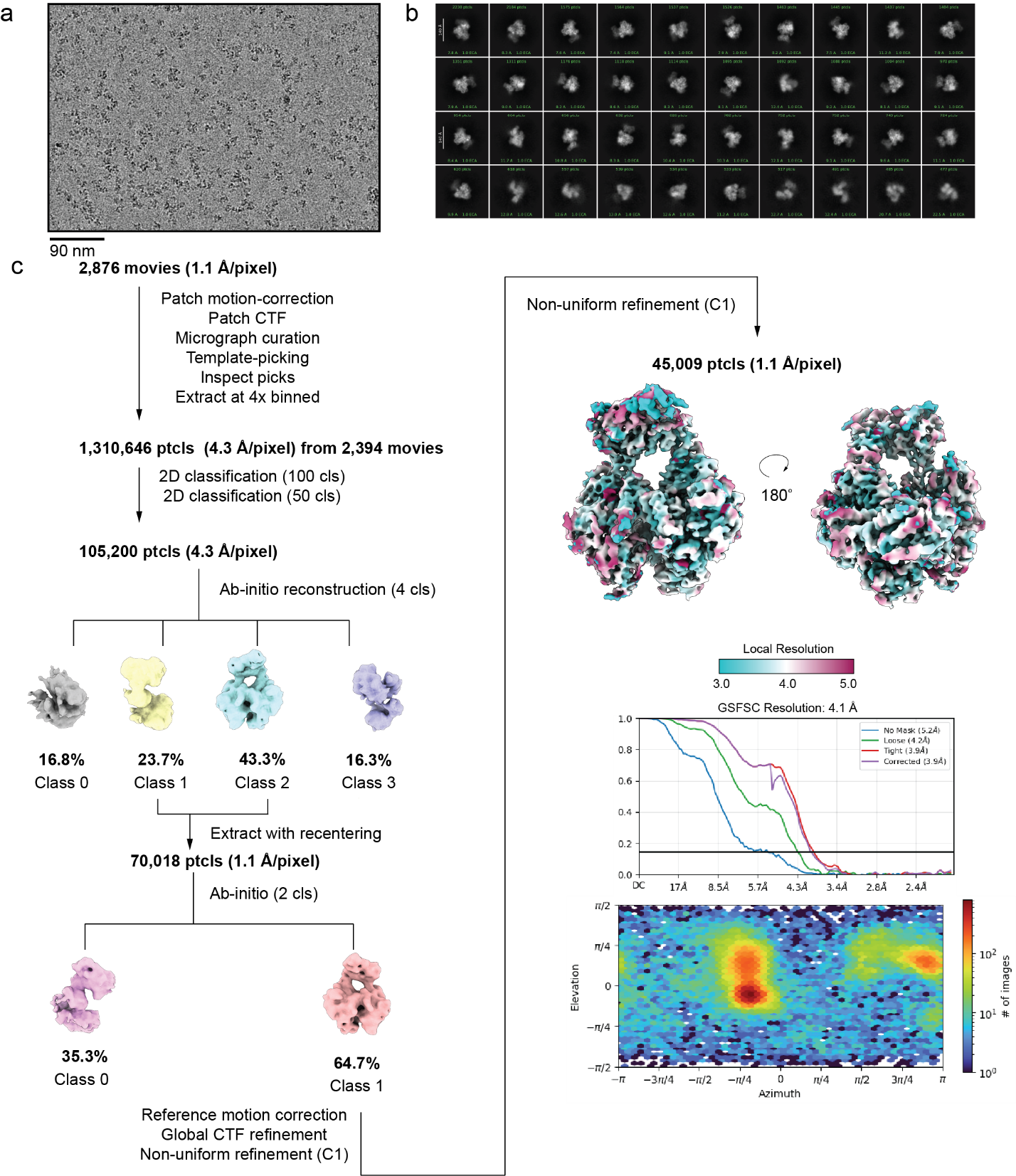

**Supplementary Figure 11 - Data processing pipeline for Polymerase α-primase and PvLEA4 dataset.** 1 µM Polα-primase with 8 µM PvLEA4. **(a)** Representative micrograph. **(b)** Top 40 2D classes. **(c)** Cryo-EM processing pipeline used to obtain final map of Polα-primase.

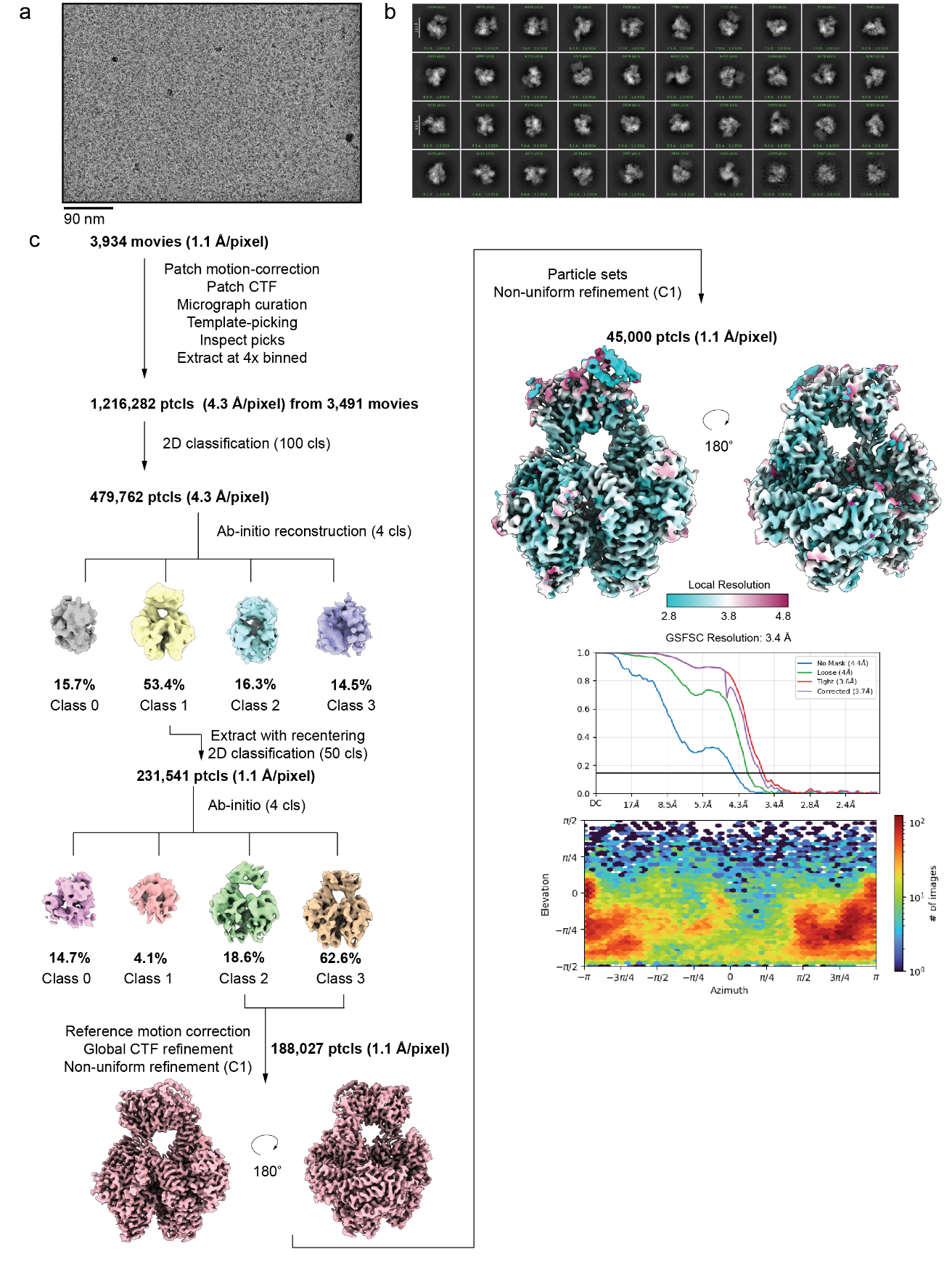

**Supplementary Figure 12 - Data processing pipeline for Polymerase α-primase and AtLEA4-5 dataset.** 1 µM Polα-primase with 8 µM AtLEA4-5. **(a)** Representative micrograph. **(b)** Top 40 2D classes. **(c)** Cryo-EM processing pipeline used to obtain final map of Polα-primase

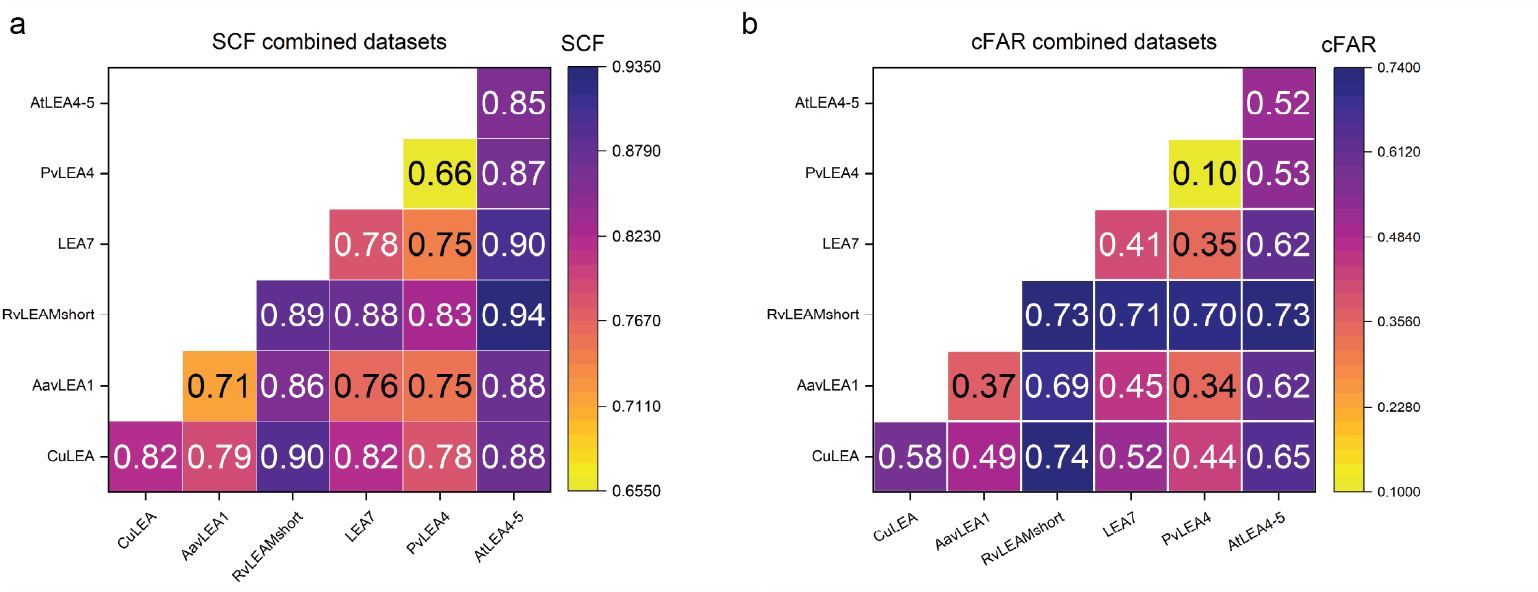

**Supplementary Figure 13 - Polymerase α-primase post-processing combined dataset orientation metric heatmap.** All values obtained from cryoSPARC. **(a)** Sample Compensation Factor (SCF) values. A lower value indicates more severe orientation bias and a value above 0.8 is generally considered good sampling. **(b)** Conical FSC Area Ratio (cFAR) values. The minimum and maximum area under the FSC curves. A value close to 0.5 indicates isotropic distribution while a small value indicates severe orientation bias.

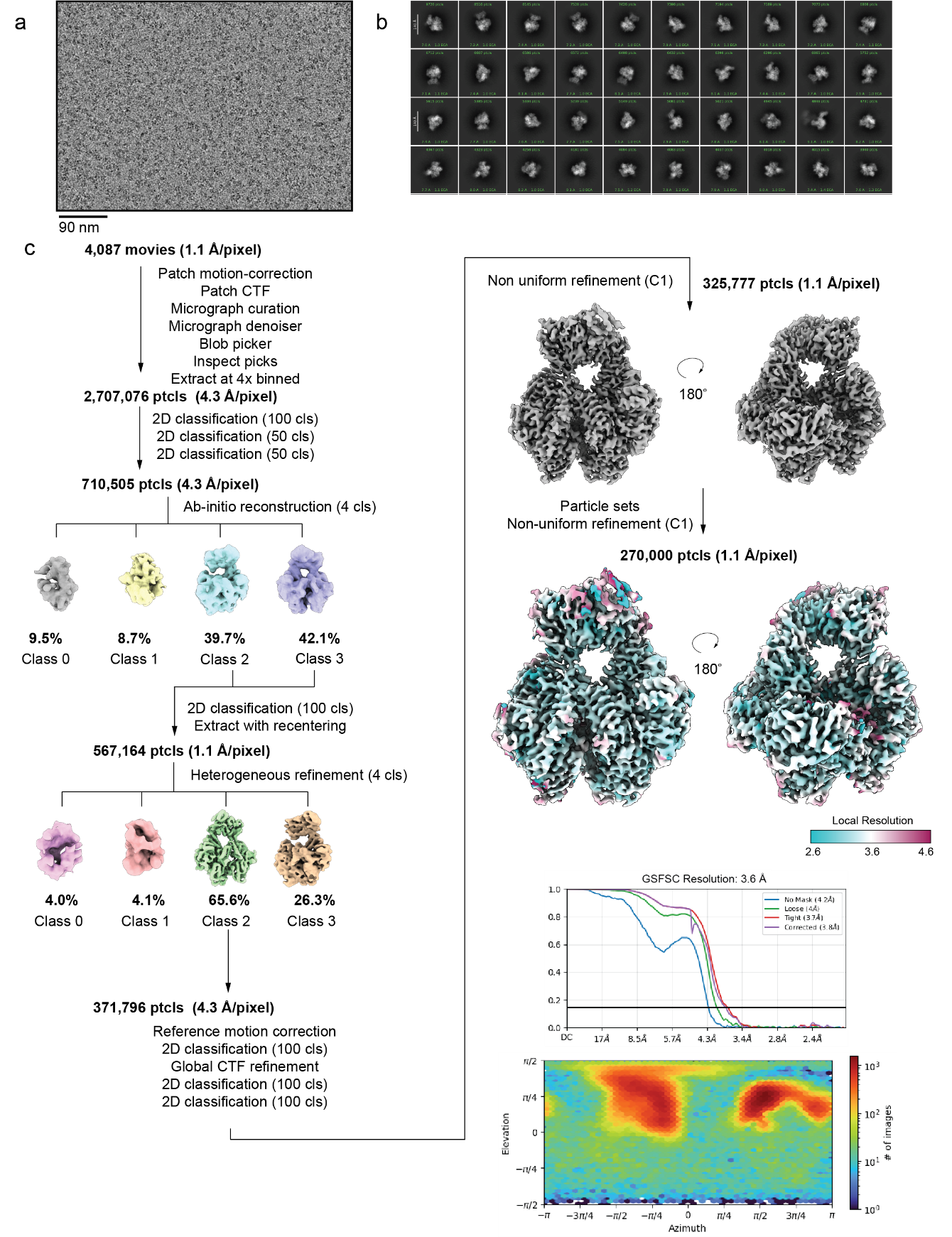

**Supplementary Figure 14 - Data processing pipeline for Polymerase α-primase and all LEAs combined dataset.** 1 µM Polα-primase with 8 µM all LEAs (~1.33 µM each). **(a)** Representative micrograph. **(b)** Top 40 2D classes. **(c)** Cryo-EM processing pipeline used to obtain final map of Polα-primase.

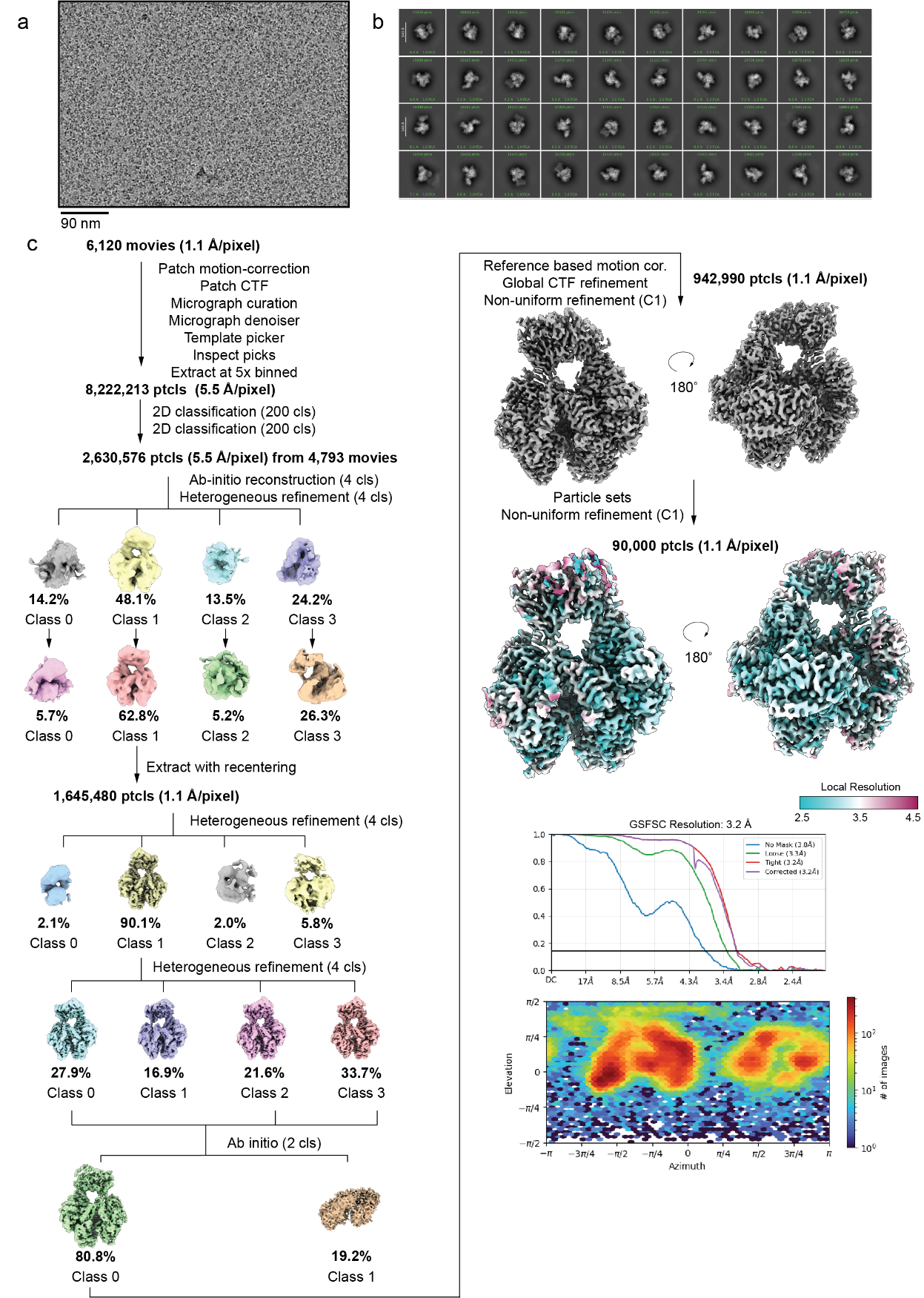

**Supplementary Figure 15 - Data processing pipeline for Polymerase α-primase and AtLEA4-5 + RvLEAM_short_.** 1 µM Polα-primase with 8 AtLEA4-5 + RvLEAM_short_ (~4 µM each). **(a)** Representative micrograph. **(b)** Top 40 2D classes. **(c)** Cryo-EM processing pipeline used to obtain final map of Polα-primase.

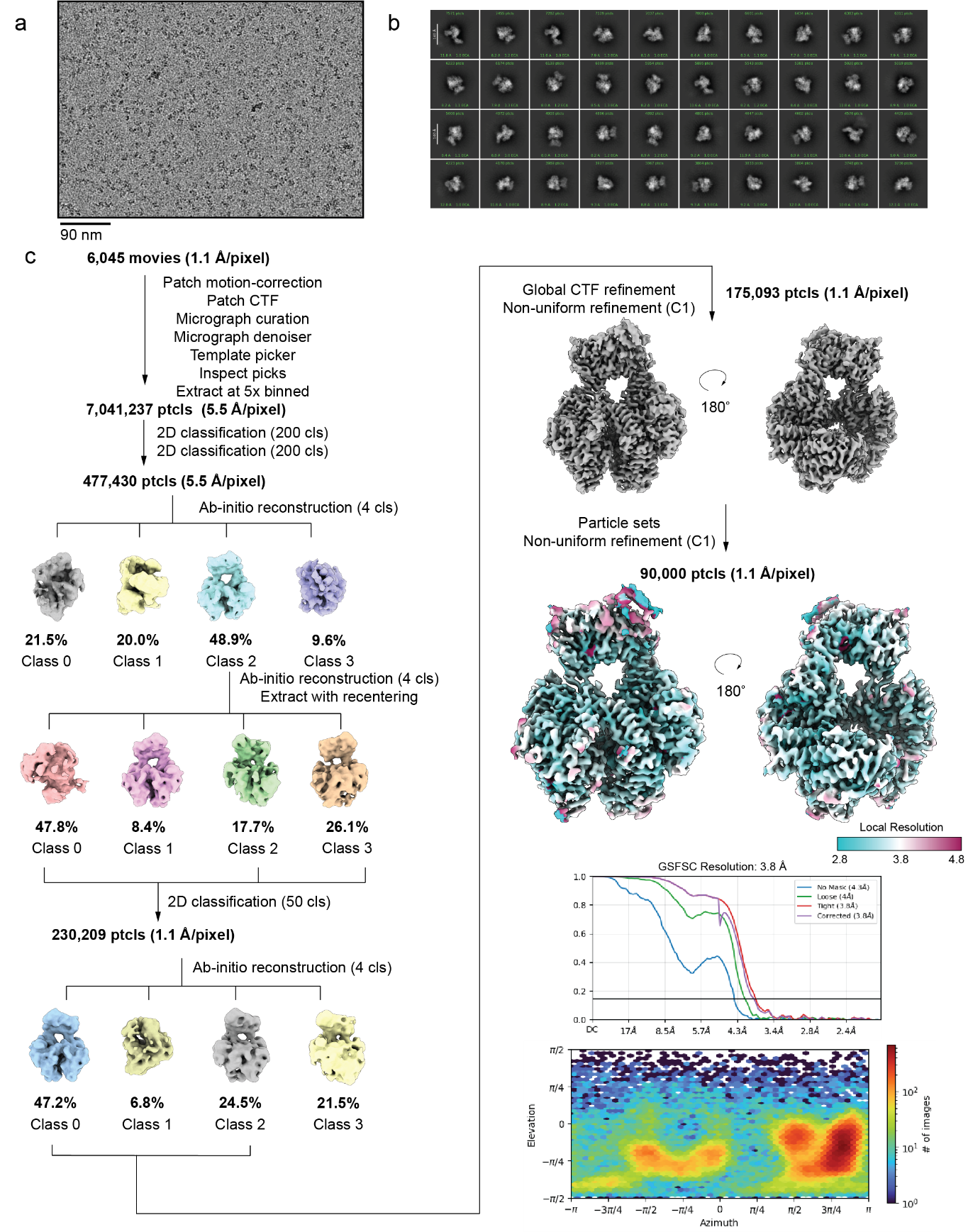

**Supplementary Figure 16 - Data processing pipeline for Polymerase α-primase and AtLEA4-5 + PvLEA4.** 1 µM Polα-primase with 8 µM AtLEA4-5 + PvLEA4 (~4 µM each). **(a)** Representative micrograph. **(b)** Top 40 2D classes. **(c)** Cryo-EM processing pipeline used to obtain final map of Polα-primase.

**Supplementary Table 1:** Orientation metrics for catalase datasets. 90,000 particles each.

|  | **cFAR** | **SCF** | **Sphericity** | **3DFSC** | **Global res.** |
| --- | --- | --- | --- | --- | --- |
| **No LEA** | 0.23 | 0.284 | 0.928 | 3.24 Å | 3.06 Å |
| **AavLEA1** | 0.49 | 0.626 | 0.963 | 2.81 Å | 2.79 Å |
| **RvLEAM_short_** | 0.59 | 0.578 | 0.961 | 3.27 Å | 3.27 Å |

**Supplementary Table 2:** Orientation metrics for HA trimer datasets. 87,122 particles each

|  | **cFAR** | **SCF** | **Sphericity** | **3DFSC** | **Global res.** |
| --- | --- | --- | --- | --- | --- |
| **No LEA** | 0.03 | 0.301 | 0.826 | 3.51 Å | 3.35 Å |
| **AavLEA1** | 0.55 | 0.918 | 0.982 | 3.70 Å | 3.68 Å |
| **RvLEAM_short_** | 0.51 | 0.838 | 0.975 | 3.58 Å | 3.55 Å |

**Supplementary Table 3:** Orientation metrics for Polα-primase datasets. 45,000 particles each.

|  | **cFAR** | **SCF** | **Sphericity** | **3DFSC** | **Global res.** |
| --- | --- | --- | --- | --- | --- |
| **CuLEA** | 0.58 | 0.818 | 0.984 | 4.10 Å | 4.03 Å |
| **AavLEA1** | 0.37 | 0.714 | 0.916 | 3.45 Å | 3.55 Å |
| **RvLEAM_short_** | 0.73 | 0.885 | 0.986 | 3.51 Å | 3.51 Å |
| **LEA7** | 0.41 | 0.781 | 0.929 | 3.58 Å | 3.58 Å |
| **PvLEA4** | 0.10 | 0.655 | 0.822 | 4.10 Å | 4.10 Å |
| **AtLEA4-5** | 0.52 | 0.853 | 0.974 | 3.83 Å | 3.83 Å |

**Supplementary Table 4:** Orientation metrics for combined LEA- Polα-primase datasets. 90,000 particles each unless notated with *.

|  |  | **cFAR** | **SCF** | **Sphericity** | **3DFSC** | **Global res.** |
| --- | --- | --- | --- | --- | --- | --- |
| **Experimental combination** | **All LEAs*** | 0.43 | 0.682 | 0.951 | 3.87 Å | 3.75 Å |
|  | **RvLEAM_short_ & AtLEA4-5** | 0.40 | 0.700 | 0.969 | 3.24 Å | 3.17 Å |
|  | **PvLEA4 & AtLEA4-5** | 0.49 | 0.725 | 0.970 | 3.96 Å | 3.82 Å |
| **Computational combination** | **All LEAs*** | 0.69 | 0.877 | 0.986 | 3.34 Å | 3.28 Å |
|  | **RvLEAM_short_ & AtLEA4-5** | 0.75 | 0.935 | 0.986 | 3.44 Å | 3.37 Å |
|  | **PvLEA4 & AtLEA4-5** | 0.51 | 0.865 | 0.979 | 3.70 Å | 3.59 Å |

*270,000 particles

**Supplementary Table 5**: Cryo-EM data collection, refinement, and validation statistics for Catalase.

|  | Catalase (1 µM) | Catalase (1 µM), AavLEA1 (8 µM) | Catalase (1 µM), RvLEAMshort (8 µM) |
| --- | --- | --- | --- |
| **Data collection and processing** | PDB-36OG  EMD-77717 | PDB-36OH  EMD-77718 | EMD-77719 |
| Magnification | 79,000 | 79,000 | 79,000 |
| Voltage (kV) | 200 | 200 | 200 |
| Electron exposure (e–/Å^2^) | 50 | 50 | 50 |
| Defocus range (μm) | -2.5 – (-1.0) | -2.5 – (-1.0) | -2.5 – (-1.0) |
| Pixel size (Å) | 1.064 | 1.064 | 1.064 |
| Symmetry imposed | D2 | D2 | D2 |
| Initial particle images (no.) | 1,986,990 | 399,852 | 1,163,787 |
| Final particle images (no.) | 90,000 | 89,816 | 90,000 |
| Map resolution (Å)  FSC threshold | 3.0 Å  0.143 | 2.8 Å  0.143 | 3.3 Å  0.143 |
| **Refinement** |  |  |  |
| Initial model used (PDB code)  Model-Map CC score | 7P8W  0.84 | 7P8W  0.81 | 7P8W  0.74 |
| Model resolution (Å)  FSC threshold | 3.3 Å  0.143 | 2.8 Å  0.143 | 3.3 Å  0.143 |
| Map sharpening *B* factor (Å^2^) | 123.4 | 104.2 | 147.6 |
| Q-Score | 0.71 | 0.73 | 0.63 |
| Model composition  Non-hydrogen atoms  Protein residues  Ligands | 17668  2008  NDP:4, HEM:4 | 17668  2008  NDP:4, HEM:4 | 17668  2008  NDP:4, HEM:4 |
| *B* factors (Å^2^)  Protein  Ligand | 50.45  45.28 | 42.03  34.65 | 59.37  53.72 |
| R.m.s. deviations  Bond lengths (Å)  Bond angles (°) | 0.003 (0)  0.587 (0) | 0.003 (0)  0.540 (0) | 0.002 (0)  0.539 (0) |
| Validation  MolProbity score  Clashscore  Poor rotamers (%) | 1.75  7.13  1.10 | 1.56  5.92  0.92 | 1.72  7.72  1.10 |
| Ramachandran plot  Favored (%)  Allowed (%)  Disallowed (%) | 95.25  4.75  0.00 | 96.40  3.60  0.00 | 96.05  3.95  0.00 |

**Supplementary Table 6:** Cryo-EM data collection, refinement and validation statistics for HA trimer.

|  | HA-trimer (1 µM) | HA-trimer (1.5 µM), AavLEA1 (12 µM) | HA-trimer (1.5 µM), RvLEAMshort (6 µM) |
| --- | --- | --- | --- |
| **Data collection and processing** | PDB-35UY  EMD-77215 | PDB-35UZ  EMD-77216 | PDB-35VA  EMD-77217 |
| Magnification | 79,000 | 79,000 | 79,000 |
| Voltage (kV) | 200 | 200 | 200 |
| Electron exposure (e–/Å^2^) | 50 | 50 | 50 |
| Defocus range (μm) | -2.5 – (-1.0) | -2.5 – (-1.0) | -2.5 – (-1.0) |
| Pixel size (Å) | 1.064 | 1.064 | 1.064 |
| Symmetry imposed | C3 | C3 | C3 |
| Initial particle images (no.) | 1,340,513 | 1,725,541 | 1,545,514 |
| Final particle images (no.) | 87,122 | 87,122 | 87,122 |
| Map resolution (Å)  FSC threshold | 3.4 Å  0.143 | 3.7 Å  0.143 | 3.6 Å  0.143 |
| **Refinement** |  |  |  |
| Initial model used (PDB code)  Model-Map CC score | 6W8B  0.67 | 6W8B  0.80 | 6W8B  0.82 |
| Model resolution (Å)  FSC threshold | 3.3 Å  0.143 | 3.6 Å  0.143 | 3.5 Å  0.143 |
| Map sharpening *B* factor (Å^2^) | 113.2 | 168.6 | 157.9 |
| Q-Score | 0.52 | 0.63 | 0.66 |
| Model composition  Non-hydrogen atoms  Protein residues  Ligands | 11799  1455  BMA:3, NAG:18,  MAN: 6 | 11799  1455  BMA:3, NAG:18,  MAN: 6 | 11799  1455  BMA:3, NAG:18,  MAN: 6 |
| *B* factors (Å^2^)  Protein  Ligand | 34.84  42.45 | 37.97  58.67 | 50.23  73.90 |
| R.m.s. deviations  Bond lengths (Å)  Bond angles (°) | 0.009 (0)  1.302 (0) | 0.004 (0)  0.708 (0) | 0.004 (0)  0.588 (0) |
| Validation  MolProbity score  Clashscore  Poor rotamers (%) | 2.22  14.73  2.20 | 1.61  8.34  0.39 | 1.67  7.17  0.31 |
| Ramachandran plot  Favored (%)  Allowed (%)  Disallowed (%) | 95.91  4.02  0.07 | 97.09  2.77  0.14 | 95.98  3.88  0.14 |

**Supplementary Table 7.** Cryo-EM data collection, refinement and validation statistics for Polymerase α-primase (PP).

|  | PP (1 µM), CuLEA (8 µM) | PP (1 µM)  RvLEAM_short_ (8 µM) | PP (1 µM) LEA7 (8 µM) | PP (1 µM) PvLEA4 (8 µM) | PP (1 µM) AtLEA4-5 (8 µM) | PP (1 µM) All LEAs (8 µM) | PP (1 µM) RvLEAM_short_ & AtLEA4-5 (8 µM) | PP (1 µM) PvLEA4& AtLEA4-5 ( 8µM) |
| --- | --- | --- | --- | --- | --- | --- | --- | --- |
| **Data collection and processing** | EMD-77626 | EMD-77627 | EMD-77628 | EMD-77629 | EMD-77625 | EMD- | EMD- | EMD- |
| Magnification | 79,000 | 79,000 | 79,000 | 79,000 | 79,000 | 79,000 | 79,000 | 79,000 |
| Voltage (kV) | 200 | 200 | 200 | 200 | 200 | 200 | 200 | 200 |
| Electron exposure (e–/Å^2^) | 50 | 50 | 50 | 50 | 50 | 50 | 50 | 50 |
| Defocus range (μm) | -2.5 – (-1.0) | -2.5 – (-1.0) | -2.5 – (-1.0) | -2.5 – (-1.0) | -2.5 – (-1.0) | -2.5 – (-1.0) | -2.5 – (-1.0) | -2.5 – (-1.0) |
| Pixel size (Å) | 1.064 | 1.064 | 1.064 | 1.064 | 1.064 | 1.064 | 1.064 | 1.064 |
| Symmetry imposed | C1 | C1 | C1 | C1 | C1 | C1 | C1 | C1 |
| Initial particle images (no.) | 2,055,454 | 2,865,601 | 2,538,992 | 1,310,646 | 1,216,282 | 2,622,992 | 8,222,213 | 7,045,237 |
| Final particle images (no.) | 45,000 | 45,000 | 45,000 | 45,009 | 45,000 | 270,000 | 90,000 | 90,000 |
| Map resolution (Å)  FSC threshold | 4.0 Å  0.143 | 3.4 Å  0.143 | 3.5 Å  0.143 | 3.9 Å  0.143 | 3.7 Å  0.143 | 3.8 Å  0.143 | 3.2 Å  0.143 | 3.8 Å  0.143 |
| Map sharpening *B* factor (Å^2^) | 119.1 | 98.3 | 99.8 | 100.8 | 111.0 | 163.3 | 100.2 | 120.4 |
